# Rapid recall and *de novo* T cell responses during SARS-CoV-2 breakthrough infection

**DOI:** 10.1101/2022.12.19.521129

**Authors:** Marios Koutsakos, Arnold Reynaldi, Wen Shi Lee, Julie Nguyen, Thakshila Amarasena, George Taiaroa, Paul Kinsella, Kwee Chin Liew, Thomas Tran, Helen E Kent, Hyon-Xhi Tan, Louise C Rowntree, Thi H O Nguyen, Paul G Thomas, Katherine Kedzierska, Jan Petersen, Jamie Rossjohn, Deborah A Williamson, David Khoury, Miles P Davenport, Stephen J Kent, Adam K Wheatley, Jennifer A Juno

**Affiliations:** Peter Doherty Institute for Infection and Immunity, Department of Microbiology and Immunology, University of Melbourne, VIC, Australia.; Kirby Institute, University of New South Wales, NSW, Australia.; Victorian Infectious Diseases Reference Laboratory, The Royal Melbourne Hospital at The Peter Doherty Institute for Infection and Immunity, VIC, Australia.; Department of Infectious Diseases, The University of Melbourne at the Peter Doherty Institute for Infection and Immunity, Melbourne, VIC 3000, Australia.; Department of Immunology, St. Jude Children’s Research Hospital, Memphis, TN, USA.; Global Station for Zoonosis Control, Global Institution for Collaborative Research and Education (GI-CoRE) Hokkaido University, Sapporo, Japan; Infection and Immunity Program and The Department of Biochemistry and Molecular Biology, Biomedicine Discovery Institute Monash University, Clayton, Victoria, Australia.; Institute of Infection and Immunity, Cardiff University School of Medicine, Heath Park, Cardiff, UK.; Melbourne Sexual Health Centre and Department of Infectious Diseases, Alfred Hospital and Central Clinical School, Monash University, Melbourne, VIC, Australia. (lead contact)

## Abstract

While the protective role of neutralising antibodies against COVID-19 is well-established, questions remain about the relative importance of cellular immunity. Using 6 pMHC-multimers in a cohort with early and frequent sampling we define the phenotype and kinetics of recalled and primary T cell responses following Delta or Omicron breakthrough infection. Recall of spike-specific CD4^+^ T cells was rapid, with cellular proliferation and extensive activation evident as early as 1 day post-symptom onset. Similarly, spike-specific CD8^+^ T cells were rapidly activated but showed variable levels of expansion. Strikingly, high levels of SARS-CoV-2-specific CD8^+^ T cell activation at baseline and peak were strongly correlated with reduced peak SARS-CoV-2 RNA levels in nasal swabs and accelerated clearance of virus. Our study demonstrates rapid and extensive recall of memory T cell populations occurs early after breakthrough infection and suggests that CD8^+^ T cells contribute to the control of viral replication in breakthrough SARS-CoV-2 infections.

## Introduction

COVID-19 vaccines provide a degree of protection against acquisition of infection but robustly protect against severe disease in the event of vaccine breakthrough infection (BTI)(Lauring et al., 2022), even in the face of antibody-evasive viral variants such as BA.1 and BA.2(Buchan et al., 2022; Kirsebom et al., 2022). Neutralising antibody titres in blood are well established as a correlate of protection from both SARS-CoV-2 infection and severe COVID-19 disease(Cromer et al., 2022; Khoury et al., 2021), with evidence for a mechanistic protective role supported by the clinical utility of monoclonal antibody treatments(Stadler et al., 2022). Nonetheless, studies in humans and animal models suggest that multiple immunological effectors likely contribute to viral control and clearance(Liu et al., 2022; Zhuang et al., 2021). In particular, it has been proposed that CD4^+^ and CD8^+^ T cell memory are important mediators of vaccine-associated protection from severe outcomes (Bertoletti et al., 2022b; Scurr et al., 2022; Wherry and Barouch, 2022). Interestingly, although unvaccinated and vaccinated cohorts show similar peak viral load after infection, vaccinated individuals exhibit accelerated viral clearance in the upper respiratory tract (URT) beginning four to six days after symptom onset(Chia et al., 2022; Garcia-Knight et al., 2022; Puhach et al., 2022; Singanayagam et al., 2022). While the mechanisms of vaccine-associated viral decline are yet to be defined, a role is plausible for both CD8^+^ T cells, which may induce cytolysis or produce antiviral cytokines (Rha et al., 2021; Sekine et al., 2020) as well as CD4^+^ T cells, which may support the recall of humoral immunity or potentially exert cytotoxic activity (Kaneko et al., 2022; Meckiff et al., 2020). There is, however, a paucity of data directly linking CD8^+^ and/or CD4^+^ T cell recall to viral clearance in human SARS-CoV-2 infections.

Previous studies of BTI have demonstrated the recall of spike (S)-specific memory T cells established by prior immunisation, as well as the induction of primary T cell responses against non-vaccine encoded viral antigens such as nucleocapsid (Collier et al., 2022; Kared et al., 2022; Lim et al., 2022; Minervina et al., 2022). To have a meaningful impact on either the rate of viral clearance or the likelihood of progressing to severe disease, memory T cell responses must be efficiently activated, likely within the first few days, following BTI(Bertoletti et al., 2022a). While cross-sectional studies of BTI cohorts have provided evidence for variable activation or expansion of S-specific T cells(Collier *et al*., 2022; Kared *et al*., 2022), early longitudinal sampling in cohorts with well-defined exposure history is rare(Kedzierska and Thomas, 2022; Koutsakos et al., 2022; Reinscheid et al., 2022). Additional work is therefore required to understand how quickly T cell activation, proliferation and effector function can occur relative to viral clearance.

In addition to the paucity of immunological studies that are temporally aligned with viral kinetics, determination of T cell activation and *ex vivo* phenotype requires direct detection of antigen-specific T cells without *in vitro* restimulation. The use of fluorescently conjugated pMHC-multimers can allow for sensitive detection of antigen-specific T cells and their ex-vivo phenotypic characterisation in cohorts with known HLA alleles(Altman et al., 1996). Indeed, the analyses of antigen-specific CD4^+^ and CD8^+^ T cells using pMHC multimers have provided novel insights into the biogenesis and maintenance of T cell responses following SARS-CoV-2 vaccination and primary infection(Jung et al., 2022; Mudd et al., 2022; Oberhardt et al., 2021; Wragg et al., 2022). Here, we use 6 pMHC-I and pMHC-II multimers presenting known immunodominant SARS-COV-2 viral epitopes(Habel et al., 2020; Minervina *et al*., 2022; Mudd *et al*., 2022; Nguyen et al., 2021; Peng et al., 2022; Rowntree et al., 2021; Wragg *et al*., 2022) to precisely define the frequency and phenotypes of SARS-CoV-2 specific CD8^+^ and CD4^+^ T cells during both the earliest events post-BTI and over extended timelines of periodic antigen re-exposure. Further, we compare the kinetics of CD8^+^ T cell recall to the induction of a primary CD8^+^ T cell response to SARS-CoV-2 nucleocapsid, a non-vaccine encoded viral antigen. These data define the relationship between viral replication in the upper respiratory tract and T cell activation, proliferation, and initiation of effector programs, shedding light on the dynamics of human adaptive immunity to respiratory virus infection in a highly vaccinated population.

## Results

### Kinetics of viral clearance and antibody recall following SARS-CoV-2 breakthrough infections

To understand the kinetics of T cell recall in relation to viral clearance and humoral immunity, we recruited a cohort of 23 individuals with PCR-confirmed SARS-CoV-2 breakthrough infection (BTI) with Delta (n= 6), Omicron BA.1 (n= 7) or Omicron BA.2 (n= 10) strains (Figure 1A, Table S1). Frequent longitudinal nasal swabs and peripheral blood samples were obtained 0-14 days post-symptom onset (PSO) with additional follow-up blood samples obtained up to day 44 PSO. Each individual provided on average 9 (range 4-13) samples with a total of 150 nasal swabs and 138 blood samples analysed. All BTIs were mild in severity and occurred on average 100 days from last vaccination (for the 21/23 participants with exact date of vaccination available).

**Figure 1.**
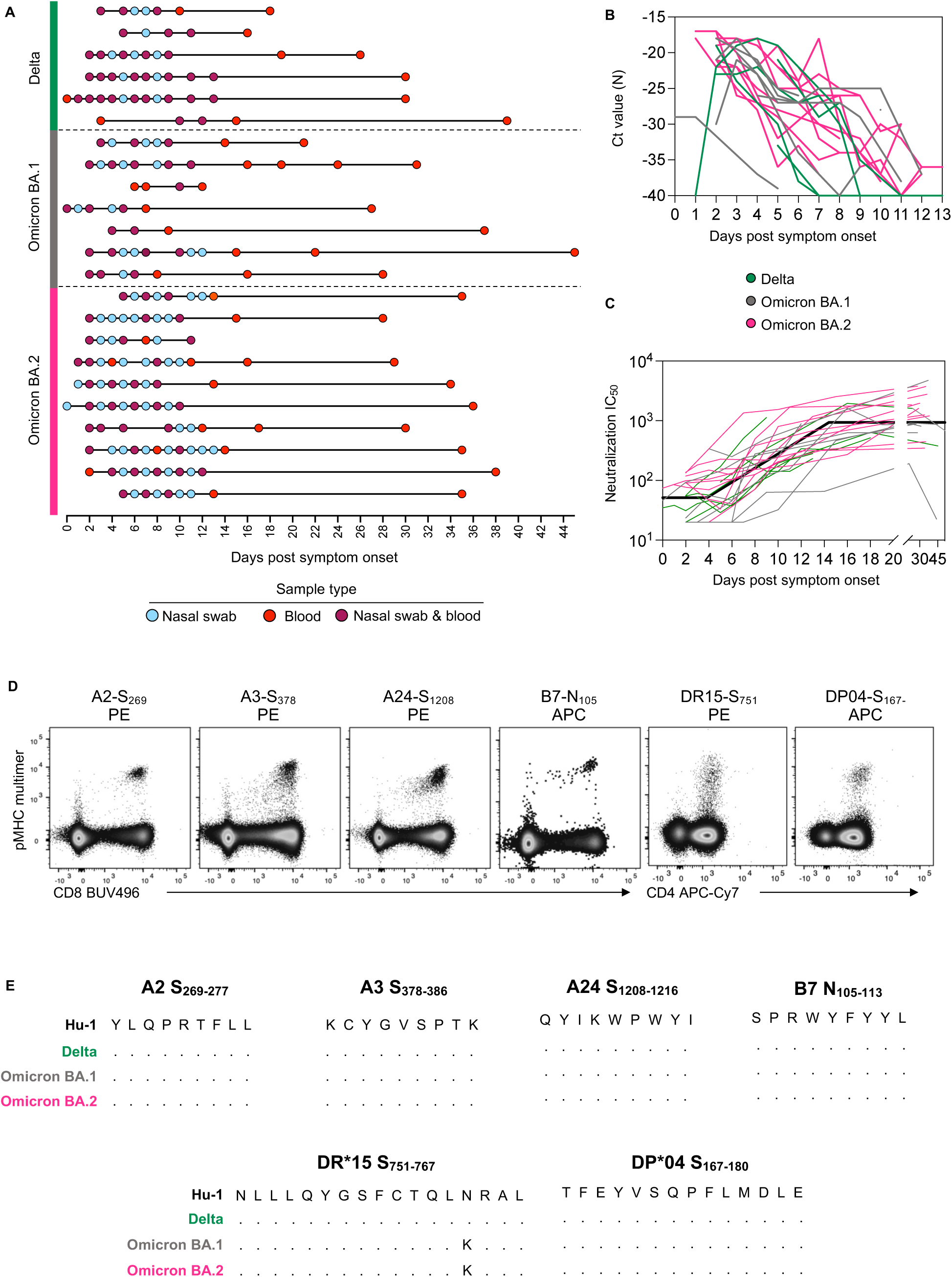
Fine longitudinal sampling of SARS-CoV-2 breakthrough infection and study design. **(A)** Schematic of longitudinal sample collection following breakthrough infection with Delta, Omicron BA.1 or Omicron BA.2. Each line represents a single donor and each point represents a sample collection (blue, nasal swab; red, blood; purple, both swab and blood). **(B)** Kinetics of viral load measured by Ct values for SARS-CoV-2 N gene in serial nasal swabs. Green, Delta breakthrough infection; grey, BA.1; pink, BA.2. **(C)** Kinetics of neutralizing antibodies measured by a live virus microneutralization assay using an antigenically similar virus to the infecting VOC. For (B) and (C) n = 6 participants for Delta, n = 7 for Omicron BA.1 and n=10 for Omicron BA.2. For (C) the bold black line represents the mean estimate from the piecewise linear regression model using the estimated parameters. **(D)** Representative flow cytometry plots for each pMHC multimer used for the detection of antigen-specific CD8^+^ and CD4^+^ T cells. Data were collected from cryopreserved PBMC following SARS-CoV-2 infection. **(E)** Sequence alignment of peptides with HLA restriction used for the detection of antigen-specific CD8^+^ and CD4^+^ T cells across selected SARS-CoV-2 VOCs.

Viral load was determined in nasal swabs by qPCR of nucleocapsid (N) gene (Figure 1B). Viral load peaked early after symptom onset (median day 3 PSO but often at the earliest sample collected), with no significant difference in peak viral load across the three VOCs. Analysis of the rate of viral clearance determined by linear regression showed significantly faster viral decay among Delta breakthroughs compared to Omicron (Supp Table 2).

The recall of neutralising antibodies was analysed using a live virus neutralisation assay (Figure 1C). The kinetics of immune recall following BTI were estimated using piecewise linear regression to estimate: (1) duration of time between symptom onset and earliest change in neutralising antibody titre (2) time of peak response relative to symptom onset, and (3) rates of growth and decay. Antibody titres against the infecting VOC (or an antigenically similar strain) increased from day 4.3 PSO (95% CI = 2.2 – 8.4) (Supp Table 2), and peaked around day 14.6 (95% CI = 12.2 – 17.4) (similar to previous reports(Koutsakos *et al*., 2022)). There were no significant differences in neutralising antibody kinetics between subjects infected with the Delta, BA.1 or BA.2 VOCs when assessed by piecewise linear regression modelling (Supp Table 2).

### Characterisation of antigen-specific T cell responses using pMHC multimers

To conduct a detailed characterisation of CD4^+^ and CD8^+^ T cell recall, we selected four CD8^+^ and two CD4^+^ T cell epitopes from SARS-CoV-2 that are restricted by HLA-alleles found at high frequency in our cohort. Three immunoprominent MHC class I restricted S derived epitopes (HLA-A*02:01-S_269-277_, HLA-A*03:01-S_378-386_, HLA-A*24:02-S_1208-1216_) and one N derived epitope (HLA-B*07:02-N_105-113_) were assessed for CD8^+^ T cells, while two MHC class II restricted S derived epitopes (HLA-DRB1*15:01-S_751-767_, HLA-DPB1*04:01-S_167-180_) were assessed for CD4^+^ T cells. All multimers have been previously verified for specificity(Mudd *et al*., 2022; Nguyen *et al*., 2021; Oberhardt *et al*., 2021; Rowntree *et al*., 2021; Wragg *et al*., 2022), with epitope-specific cells readily detected *ex vivo* in samples from infected individuals (Figure 1D, gating in Supp Fig 1). All peptides are well conserved across the relevant infecting VOCs in this study, with only one mutation (N764K) found for the DR15-restricted S_751_ peptide in BA.1 and BA.2 sequences (Figure 1E), which is not predicted to alter MHC binding (predicted affinity 97.06nM vs 80.03nM, NetMHCIIPan-3.2(Jensen et al., 2018)).

### Robust expansion and activation of S-specific CD4^+^ T cells after breakthrough infection

S-specific memory CD4^+^ T cells were detected using the DR15-S_751_- and DP04-S_167_ pMHC multimers in all participants carrying the relevant HLA alleles (Figure 2A). Longitudinal tracking showed a clear expansion of the number of S-specific CD4^+^ T cells following BTI. Analysis of T cell kinetics with piecewise linear model estimated that frequencies of S-specific CD4^+^ T cells rise at 2.5 days PSO (95% CI = 1.9-3.4), peaking on day 5.4 (95% CI =4.8-6.1), and slowly contracting thereafter (Figure 2B-C, Supp Table 3). Strikingly, the appearance of activated (CD38^+^ICOS^+^) Tet^+^ cells in the circulation occurred in a synchronised manner across the cohort (Figure 2A-C) with evidence of activation estimated to be initiated around 1.1 days PSO (95% CI =0.7-1.8) and peaking by day 3.6 (95% CI =3.2-4.01). Over this period the proportion of activated pMHC-II^+^ CD4^+^ T cells rose from 2.44% (95% CI =1.1-5.4) at initial sampling to a peak of 74.5% (95% CI = 68 – 81%)(Supp Table 3). Other markers of activation including CD71 and PD-1 were similarly upregulated following BTI (Figure 2D), with maximal expression between 1.6 and 4.0 days PSO (Supp Table 4). The kinetics of expansion and activation were broadly comparable between the two MHC-II epitopes, with only minor differences in the decay kinetics of some markers Supp Table 4).

**Figure 2.**
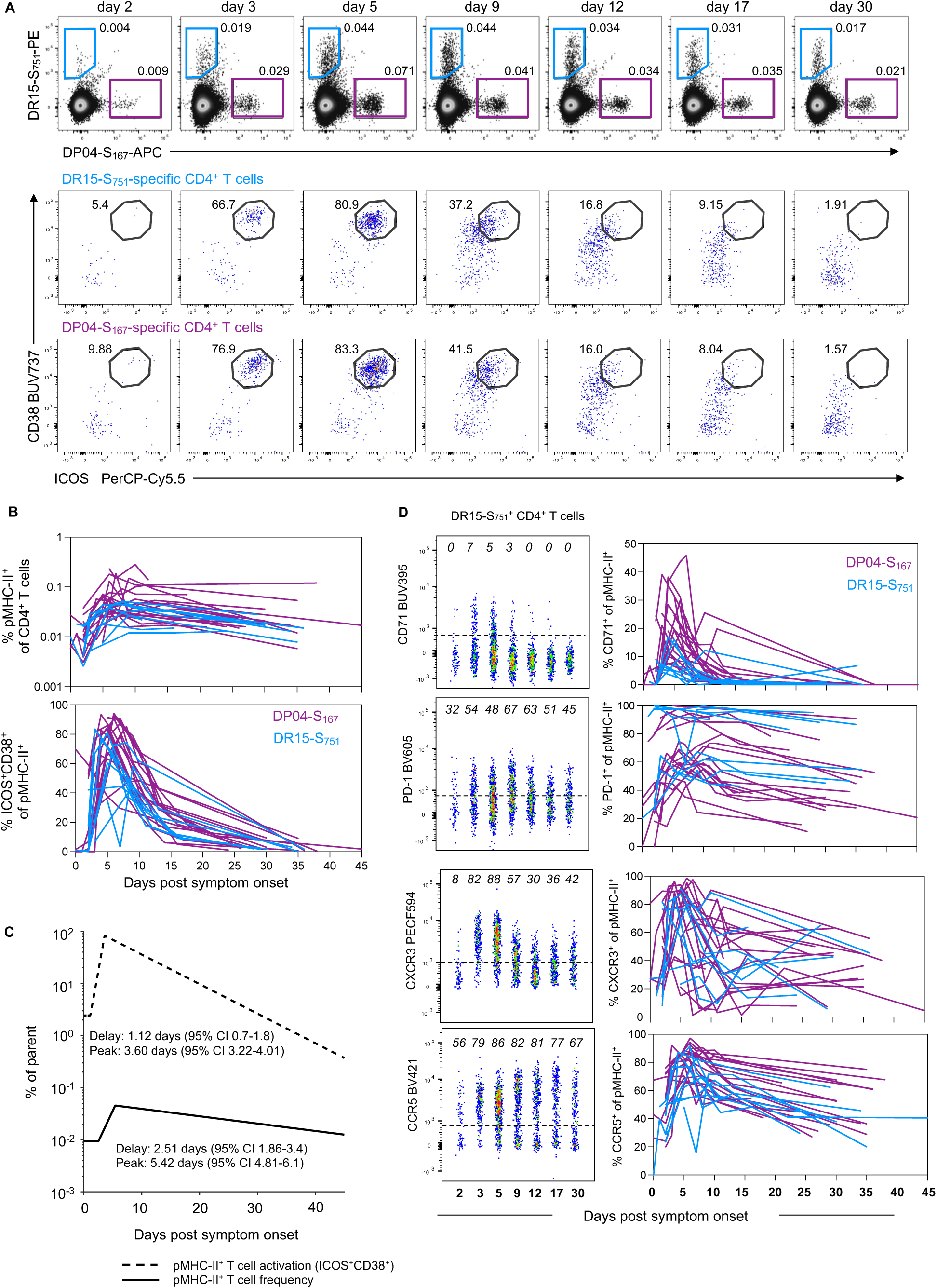
Robust expansion and activation of spike-specific CD4^+^ T cells after breakthrough infection. **(A)** Representative flow cytometry plots and kinetics of HLA-DR*15-S_751_ and HLA-DP*04-S_167_- specific CD4^+^ T cells from a single participant and co-expression of ICOS and CD38. **(B)** Representative flow cytometry plots of HLA-DR*15-S_751_ -specific CD4^+^ T cells and kinetics of phenotypic markers for both pMHC-II populations, n=19 for DP*04-S_167_ and n=9 for DR*15-S_751_. **(C)** Estimated kinetics of pMHC-II^+^ CD4^+^ T cell frequency and activated (CD38^+^ICOS^+^) phenotype. The lines indicate the mean estimate for measurement from the piecewise linear regression model, using pooled data from both pMHC-II populations (as no significant differences were found between the two). **(D)** Representative flow cytometry plots of phenotypic markers for DR*15-S_751_ (left) and kinetics marker expression for both pMHC-II populations (right). Throughout the figure, coloured lines represent individual donors for each pMHC-II-specific population, n=19 for DP*04-S_167_ and n=9 for DR*15-S_751_.

Previous studies of DR15-S_751_ and DP04-S_167_ responses demonstrated recruitment into both conventional CD4^+^ memory T cell pools and circulating T follicular helper (cTFH) cell pools following primary vaccination or infection(Mudd *et al*., 2022; Wragg *et al*., 2022). Surprisingly, we found minimal evidence of recalled T cells exhibiting a cTFH phenotype during BTI (Supp Figure 2A-B), even among individuals where CXCR5^+^ S-specific cells were clearly detected after primary vaccination. Accordingly, we found no correlation between the peak or growth rate of neutralising antibody titres and the kinetics of S-specific CD4^+^ T cell recall/memory across the cohort (Supp Figure 2C).

Phenotypically, both DR15-S_751_- and DP04-S_167_-specific cells exhibited a mixture of T_CM_-like (CD45RA^-^CCR7^+^) and T_EM_-like (CD45RA^-^CCR7^-^) profiles, with a transient increase in the relative proportion of CCR7^+^ cells occurring around the peak of T cell proliferation (Supp Figure 3A). Previously, nasopharyngeal swabs collected during mild/moderate COVID-19 revealed the upregulation of multiple inflammatory mediators, including the CXCR3 ligand CXCL10(Rajagopala et al., 2022), which may facilitate trafficking of activated T cells to the lung or nasal mucosa. S-specific CD4^+^ T cells showed upregulation of both CXCR3 and CCR5 during BTI, suggesting the potential for these cells to migrate to inflamed tissues (Figure 2D, Supp Figure 3B-C). We further observed a modest increase in granzyme B (GzmB) expression among S-specific CD4^+^ T cells by day 2 PSO (Supp Figure 3D), in line with previous reports of *GZMB* upregulation following *in vitro* culture of S-specific CD4^+^ T cells(Dong et al., 2022). Overall, we find rapid activation and expansion of S-specific CD4^+^ T cells following BTI, characterised by transient upregulation of tissue/lymphatic-homing markers.

**Figure 3.**
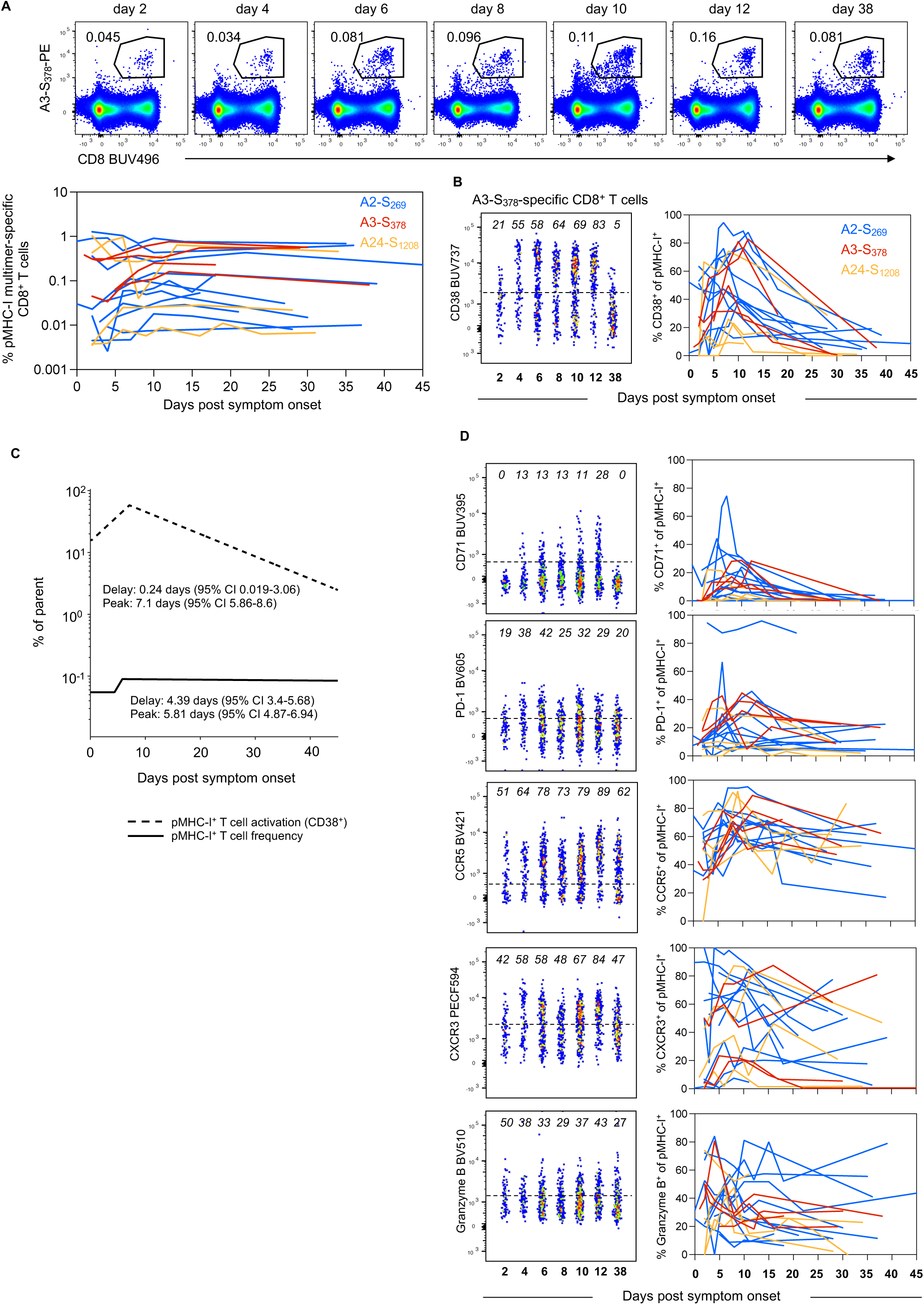
Early but variable recall of spike-specific CD8^+^ T cells after breakthrough infection. **(A)** Representative flow cytometry plots for HLA-A*3-S_378_ and kinetics of HLA-A*02-S_269_, HLA- A*03-S_378_ and HLA-A*24-S_1208_ -specific CD8^+^ T cells. **(B)** Representative flow cytometry plots for HLA-A*3-S_378_ and kinetics of activated (CD38^+^) cells for HLA-A*02-S_269_, HLA-A*03-S_378_ and HLA- A*24-S_1208_ -specific CD8^+^ T cells. (C) Estimated kinetics of pMHC-I^+^ CD8^+^ T cell frequency and activated (CD38^+^) phenotype. The lines indicate the mean estimate for measurement from the piecewise linear regression model, using pooled data from all 3 pMHC populations. **(D)** Representative flow cytometry plots of phenotypic markers for HLA-A*3-S_378_and kinetics marker expression for all 3 pMHC-I populations. Throughout the figure, coloured lines represent individual donors for each pMHC-I-specific population, n=11 for A*02-S_269_, n=4 A*03-S_378_ and n=4 for A*24- S_1208_.

### Variable expansion but universal activation of S-specific CD8^+^ T cells after breakthrough infection

S-specific CD8^+^ T cells, regardless of epitope specificity, expanded during BTI from day 4.4 PSO (95% CI = 3.4-5.7), peaking on day 5.8 (95% CI =4.9-6.9) and maintaining relatively stable frequencies thereafter (Figure 3A, Supp Figure 4A, Supp Table 5). We note, however, the variability in recall between individuals, with a subset of participants (12/18) showing considerable expansion in the number of S-specific CD8^+^ T cells (median 3.6-fold peak expansion relative to earliest timepoint available). The remaining six participants had comparatively stable frequencies of spike-specific cells over the course of follow-up (median 1.1-fold peak expansion relative to earliest timepoint available; Supp Figure 4B). Participants with limited CD8^+^ T cell expansion were significantly more likely to have received 3 vaccines compared to the rest of the cohort (p=0.038, Supp Figure 4C), and exhibited significantly higher frequencies of S-specific CD8^+^ T cells at the earliest timepoint available (p=0.017, Supp Figure 4D). There was not, however, any difference in the time interval from last vaccination to BTI between the two groups (median 74 vs 84 days, p>0.05, Supp Figure 4E).

**Figure 4.**
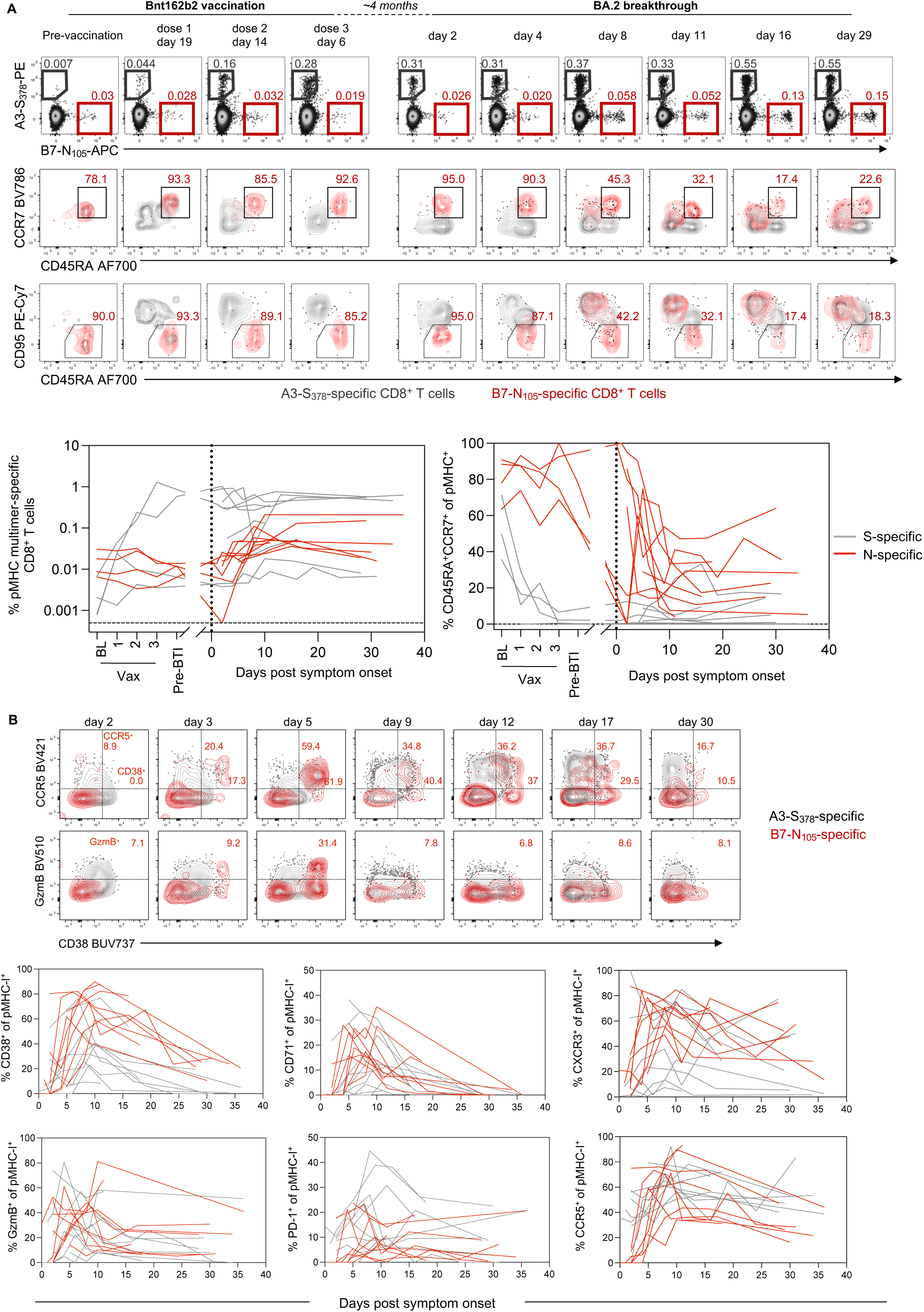
Early expansion of primary nucleocapsid-specific CD8^+^ T cells and recall of spike-specific CD8^+^ T cells after breakthrough infection. **(A)** Representative flow cytometry plots of HLA-A*02-S_269_ and HLA-B*07-N_105_-specific CD8^+^ T cells and their phenotypic analysis based on CCR7, CD45RA and CD95 from baseline throughout vaccination and subsequent SARS-CoV-2 breakthrough infection. Kinetics are shown for n=9 donors with paired analysis spike-specific CD8^+^ T cells (either A*02, A*03 and A*24) and B*07-N_105,_ for 4 of which pre-breakthrough samples were available. **(B)** Representative flow cytometry plots for HLA-A*3-S_378_ and HLA-B*07-N_105_-specific CD8^+^ T cells. Frequencies of single marker^+^ cells are shown in red for B7-N105 only. Kinetics of phenotypic markers for S- or N-specific CD8^+^ T cells following SARS-CoV-2 breakthrough infection, n=9 donors with paired spike-specific CD8^+^ T cells (either A*02, A*03 and A*24) and B*07-N_105_. Throughout the figure, coloured lines represent individual donors for each pMHC-specific population.

Despite the variable changes in T cell frequency among individuals, we observed universal upregulation of CD38 on S-specific CD8^+^ T cells, estimated to occur soon after the onset of symptoms (0.24 days PSO) and peaking at 7.1 days (Figure 3B, C). All six individuals with limited T cell expansion exhibited activation of S-specific cells, with 3 of those participants showing relatively high frequencies of CD38^+^ cells at their initial sample that declined during follow-up (Supp Figure 4F). It is therefore possible that the onset of CD8^+^ T cell activation and proliferation occurred prior to the first available sample in these individuals (i.e. before day 2 PSO), either due to a lag between infection and symptom onset, or rapid T cell recall.

In addition to CD38, we observed similar upregulation of activation marker PD-1 and proliferation marker CD71, which peaked on day 3.3 (95% CI =2.1-5.1) and day 5.4 (95% CI =4.3-6.8) PSO, respectively (Figure 3D, Supp Table 6). Consistent with our observations of S-specific CD4^+^ T cells, BTI drove upregulation of CCR5 and CXCR3 on CD8^+^ T cells (Figure 3D, Supp Figure 5A-B). During the course of infection, S-specific CD8^+^ T cells transiently changed from a CCR7^-^CD45RA^+^ to a CCR7^-/+^CD45RA^-^ phenotype (Supp Figure 5C). The expression of granzyme B was dynamic, with clearly detectible GzmB^+^ populations at the earliest timepoints post-BTI and considerable upregulation observed in some donors (Figure 3D, Supp Figure 5A). We did not note any major differences in relative immunodominance, recall kinetics or phenotype between the three S-derived pMHC-I epitopes, although the current study is not powered for a formal comparative analysis. Overall, despite variability between donors in the expansion in the number of S-specific CD8^+^ T cells, these cells consistently show high levels of activation at early timepoints after BTI.

**Figure 5.**
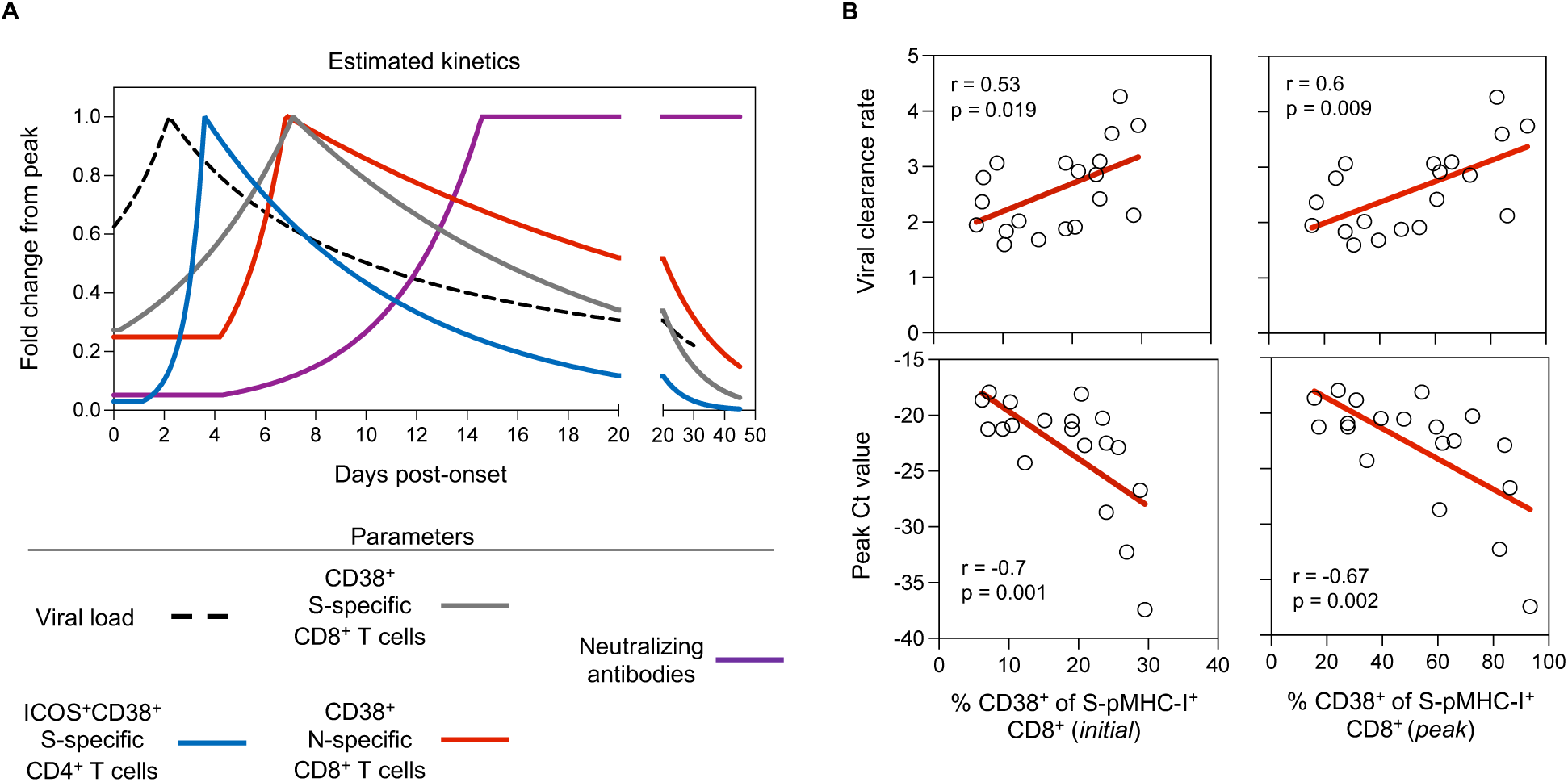
S-specific CD8^+^ T cell activation correlates with viral clearance. **(A)** Summary of estimated kinetics viral clearance and of relevant immunological parameters determined in this study**. (B)** Correlations between the initial or peak frequency of CD38^+^ S-pMHC-I^+^ CD8^+^ T cells and viral clearance rate or peak Ct value (amongst available timepoints). Spearman correlation coefficient and p-values along with a linear regression line are shown for statistically significant comparisons (p<0.05), n=19.

### Concomitant expansion of primary N-specific CD8^+^ T cells and recall of S-specific CD8^+^ T cells after breakthrough infection

All individuals in the BTI cohort were previously immunised with SARS-CoV-2 vaccines encoding only the S antigen (BNT162b2, ChAdOx-nCoV19 or mRNA-1273). Consequently, recall of vaccine-induced immunological memory targeting S could be compared to the primary immune response generated against other viral antigens. Among the 23 individuals in the BTI cohort, 9 carried both an S peptide-(A2-S_269-277_, A3-S_378_, or A24-S_1208_) and N peptide-(B7-N_105_) restricting HLA allele. Furthermore, longitudinal PBMC samples were available from four of these individuals over the course of primary vaccination, allowing us to characterise the nature of S- and N-specific CD8^+^ T cell populations prior to their BTI.

Analysis of pre-BTI samples validated previous observations of low baseline frequencies of naïve (CD45RA^+^CCR7^+^CD95^-^) S-specific CD8^+^ T cells, which increase in frequency and acquire a CD45RA^-^CCR7^+/-^CD95^+^ phenotype following vaccination (Figure 4A). At baseline, B7-N_105_-specific CD8^+^ T cells similarly presented with a naïve-like phenotype (CD45RA^+^CCR7^+^) which lacked expression of the stem-cell memory marker CD95. The frequency and naïve phenotype of these cells was stable throughout vaccination (Figure 4A), consistent with the lack of N antigen in the vaccine formulations received by these individuals. These data align with previous observations that in SARS-CoV-2 naïve donors the majority of B7-N_105_-specific CD8^+^ T cells detected *ex vivo* are antigen-inexperienced, with no evidence of priming by cross-reactive human coronavirus epitopes(Lineburg et al., 2021; Nguyen *et al*., 2021).

N-specific cells were also found at low frequency and with a naïve phenotype at early timepoints post-BTI among the wider cohort. Memory S-specific CD8^+^ T cells were ∼8 times greater in frequency than naïve N-specific CD8^+^ T cells at the first available timepoint, despite the high precursor frequency of naïve B7-N_105_ T cells observed in this and other studies(Lineburg *et al*., 2021; Nguyen *et al*., 2021) (Figure 4A). Over the course of BTI, we observed the progressive differentiation of B7-N_105_-specific T cells as they proliferated, downregulated CD45RA and acquired expression of CD95 (Figure 4A). Initiation of an effector program among N-specific CD8^+^ T cells was characterised by the concurrent upregulation of CD38, CCR5, CXCR3, GzmB and CD71 (Figure 4B). Surprisingly, however, despite the prior exposure to S-antigen and the difference in overall frequency of S- and N-specific cells, the activation kinetics of S- and N-specific CD8^+^ T cells were largely similar, with both populations exhibiting similar delay, growth rate and peak time of CD38, CD71, CXCR3, CCR5 and GzmB expression (Supp Table 7-8). Nonetheless, at early timepoints, S-specific cells expressed significantly higher levels of GzmB and CCR5 than naïve N-specific cells (Supp Table 8), providing them with the immediate effector and trafficking potential classically ascribed to memory T cells. Despite the similar kinetics, vaccine-primed S-specific CD8^+^ T cells were present at a higher frequency than N-specific CD8^+^ T cells throughout BTI. This was in contrast to a primary SARS-CoV-2 infection, where B7-N_105_-specific CD8^+^ T cells were shown to be numerically dominant population over the S-specific populations included in our analysis (Supp Fig 5E). Overall, we find both recall and primary CD8^+^ T cell responses occur at early timepoints after symptom-onset, confirming that de novo T cell responses to infection are not impaired in vaccinated individuals(Minervina *et al*., 2022).

### S-specific CD8^+^ T cell activation correlates with viral clearance

Studies of SARS-CoV-2 dynamics in the URT have demonstrated that while peak viral load is similar between vaccinated and unvaccinated individuals, prior vaccination is associated with accelerated viral clearance starting between days 4-6 PSO(Chia *et al*., 2022; Garcia-Knight *et al*., 2022; Puhach *et al*., 2022; Singanayagam *et al*., 2022). To understand the relationship between viral decline and the onset of humoral and cellular recall, we compared the kinetics of nasal viral load, CD4^+^ and CD8^+^ T cell activation, and neutralising antibody titres. Our kinetic analysis clearly indicated that activation of SARS-CoV-2 specific T cells occurs concurrently to viral clearance, and considerably earlier than recall of neutralising antibodies (Figure 5A). To further define the potential role of T cell recall in viral clearance, we performed an exploratory analysis to understand whether T cell recall kinetics correlate with levels and rate of viral clearance in the URT.

We investigated the relationship between viral dynamics (peak viral load and rate of viral clearance) with either the total frequency of antigen-specific CD4^+^ and CD8^+^ T cells or the activation of antigen-specific populations. The frequency of S-specific CD8^+^ T cells did not correlate with viral load or viral clearance rate (Supp Figure 6A). Notably, however, we identified a relationship between of S-specific CD8^+^ T cell activation and viral clearance. Specifically, the frequency of CD38^+^ S-specific cells at the time of symptom onset and at peak response both positively correlated with the rate of viral clearance (p=0.019 and 0.009, respectively, Figure 5B). Conversely, both measures of S-specific T cell activation were inversely associated with peak viral load (p=0.001 and 0.002 for initial and peak CD8^+^ T cell activation, respectively; Figure 5B). Neither the frequency nor activation of the primary N-specific CD8^+^ T cell response was significantly associated with viral peak or clearance (Supp Figure 6B-C), although this analysis is limited due to a small sample size. Additionally, we did not observe a correlation between the frequency or activation of S-specific CD4^+^ T cells and viral peak or clearance (Supp Figure 6D-E).

**Figure 6.**
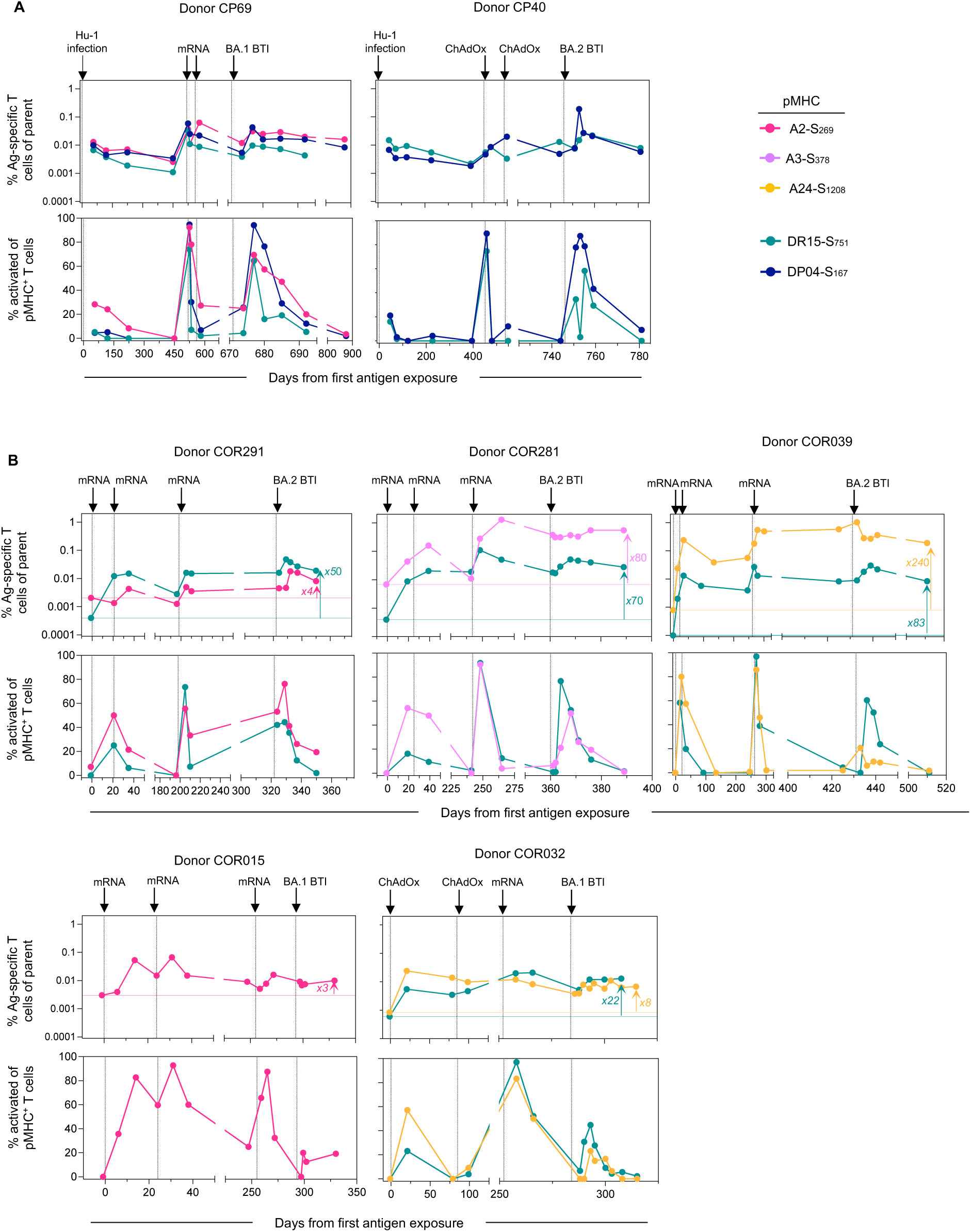
Long**-**term stability of T cell responses following multiple antigen exposures. **(A)** Kinetics for two participants with an initial Hu-1 infection. **(B)** Kinetics for 5 participants after vaccination, with no prior exposure. The horizontal line indicates the pre-exposure levels for each epitope-specific T cell population. Throughout the figure, each plot represents longitudinal data from one donor, with the frequency of pMHC^+^ cells within CD4^+^ or CD8^+^ T cells (top) and the frequency of activated (CD38^+^ICOS^+^ for pMHC-II^+^CD4^+^ T cells or CD38^+^ for pMHC-I^+^CD8^+^ T cells) cells within pMHC^+^ cells (bottom). Each arrow on the top of the plot indicates an exposure to spike antigen by infection or vaccination.

The association between S-specific CD8^+^ T cell activation and viral kinetics is intriguing, suggesting that a greater magnitude of S-specific T cell recall may be associated with a more rapid clearance of virus in the URT. However, studies of monoclonal antibody administration indicate that antibody titres may also accelerate viral clearance(Weinreich et al., 2021). Therefore, we explored whether neutralising antibody levels (to the infecting or antigenically similar strain) was also associated with viral kinetics in this cohort. We found no significant association between initial or peak neutralising antibody levels and viral peak or clearance rate in this cohort (Supp Figure 6F). Overall, the association between activation of recalled S-specific CD8^+^ T cells in blood and viral kinetics suggest that T cell activation may be an informative correlate of viral clearance following BTI.

### Long**-**term stability of T cell responses following multiple antigen exposures

As the SARS-CoV-2 pandemic progresses, the lived experience of many individuals will include recurrent exposure to the viral S protein through a combination of vaccination and infection. Seven individuals in our BTI cohort were longitudinally sampled for up to 875 days over the course of the pandemic, providing a novel insight into the long-term maintenance and recall of S-specific T cell immunity (Figure 6). This included two individuals infected with ancestral (Hu-1) virus, subsequently immunised twice with S-encoding vaccines and then acquiring an Omicron BTI (Figure 6A), and five individuals who received three doses of S-encoding vaccines and subsequently infected with Omicron (Figure 6B). While each exposure had a variable impact on the frequency of S-specific T cells, the cumulative impact of multiple vaccinations/infections was the durable maintenance of memory T cell populations substantially above pre-vaccination levels (Figure 6B). At the end of follow-up, S-specific CD8^+^ T cell frequencies were 3 to 240-fold higher compared to pre-vaccination baseline samples, with S-specific CD4^+^ T cells 22 to 83-fold higher. Notably, two participants with high S-specific T cell frequencies and limited CD8^+^ T cell expansion following BTI (COR039 and COR281) exhibited particularly high frequencies of S-specific CD8^+^ T cell following the 3^rd^ vaccine dose, comprising 0.6-1.3% of the CD8+ T cell compartment at peak (Figure 6B). Across the cohort, each exposure to S also resulted in striking but transient activation of both CD4^+^ (ICOS^+^CD38^+^) and CD8^+^ (CD38^+^) S-specific T cells. Although the sample size limits formal comparisons, breakthrough infection with BA.1 or BA.2 virus did not appear to boost circulating S-specific T cell frequencies, or their activation phenotype to a as great an extent as observed after the 3^rd^ vaccine dose (Figure 6A-B). Taken together, our longitudinal follow-up over an extended period of time demonstrates the durability and long-term maintenance of circulating vaccine-induced S-specific CD4^+^ and CD8^+^ T memory cells.

## Discussion

As SARS-CoV-2 variants increasingly escape neutralizing antibody responses, T cell responses that target conserved epitopes are likely to be of increasing immunological importance(Wherry and Barouch, 2022). Here, we characterised multiple epitope-specific CD4^+^ and CD8^+^ T cells in a longitudinal cohort of finely sampled individuals early after BTI. Our kinetic analyses clearly demonstrate rapid activation of both CD4^+^ and CD8^+^ T cells that precedes serological increases in neutralizing antibodies. Further, we find an association between high levels of S-specific CD8^+^ T cell activation and more rapid viral decline in SARS-CoV-2 RNA levels in the URT.

While mechanistic roles have been established for neutralizing antibodies in protection from acquisition of infection and progression to severe disease(Khoury *et al*., 2021; Stadler *et al*., 2022), questions have persisted over the contributions of CD4^+^ or CD8^+^ T cells (Kent et al., 2022). Identification of immune correlates that predict avoidance of severe outcomes following BTI is challenging due to the relatively low frequency of severe infections. For this reason, studies of the correlation between T cell responses and viral kinetics have been proposed(Kent *et al*., 2022) . It is pertinent to note that in mild/moderate BTI, vaccination has no apparent impact on peak viral load(Chia *et al*., 2022; Garcia-Knight *et al*., 2022; Singanayagam *et al*., 2022), with SARS-CoV-2 viral replication being curtailed at or prior to the onset of symptoms in most individuals(Koutsakos *et al*., 2022). Nonetheless, the significantly faster decline of viral load in the upper respiratory tract among vaccinated individuals in the first week post-onset(Chia *et al*., 2022; Garcia-Knight *et al*., 2022; Puhach *et al*., 2022; Singanayagam *et al*., 2022) suggests that vaccine-induced immune memory contributes to viral clearance. Our study therefore places the activation of S-specific CD4^+^ and CD8^+^ memory T cells in an immunologically relevant window following BTI.

Further support for a protective role for T cells comes from our observation that higher CD8^+^ T cell activation level in peripheral blood was associated with lower peak viral load and a faster rate of viral clearance in the URT. Importantly, being based on the estimated kinetic parameters of viral load and T cell immunity this association does not simply reflect temporal relationships between an activated immune response and declining virus. Our findings are also consistent with animal studies in which CD8^+^ T cell depletion results in delayed viral clearance(Liu *et al*., 2022). While data derived from peripheral blood may not necessarily reflect T cell responses in tissues, we find that activated S-specific T cells express chemokine receptors that should facilitate trafficking to inflamed tissues, as well as effector molecules such as GzmB that can lead to clearance of infected cells. It is important to note that we were unable to assess the role of T cell activation in the context of diverse clinical outcomes, which will need to be addressed in future studies. A key issue that remains poorly understood is whether or how accelerated viral clearance in the URT is related to protection from severe disease among vaccinated individuals. It is possible that immune responses contributing to viral clearance in the URT also prevent spread of virus to the lower respiratory tract and thus mitigate disease progression(Wherry and Barouch, 2022); in this context, both antibodies and cytolytic T cells could contribute to the containment of viral replication.

Parallel tracking of CD4^+^ and CD8^+^ T cell responses revealed the extent to which memory T cell pools can be reactivated following antigen exposure. In particular, the rapid appearance of highly activated S-specific CD4^+^ T cells in the periphery (often within a 24 hour sampling window) demonstrated the remarkable efficiency of CD4^+^ T cell recall during BTI. Furthermore, long-term tracking of S-specific T cell populations through multiple vaccinations and infections demonstrated the ability of memory cells to be established and recalled multiple times over the course of one to two years with no evidence for progressive exhaustion or anergy. There may, however, be a ‘ceiling’ of epitope-specific T cell frequencies, such that repeated antigen exposure largely maintains a stable population of memory T cells rather than progressively increasing the memory pool with each restimulation.

Previous characterization of nucleoprotein-(B7-N_105_-) specific CD8^+^ T cells in pre-pandemic samples and uninfected individuals has indicated that the majority of these cells have a naïve phenotype(Nguyen *et al*., 2021; Peng *et al*., 2022), which is further supported by our analysis. Thus, the analysis of Spike- and Nucleoprotein-specific CD8^+^ T cells in our S-vaccinated BTI cohort provides an opportunity to analyse both primary and recall CD8^+^ T cell responses together. Intriguingly, we did not find any major differences in the kinetics of primary and recalled responses, consistent with a previous report on recalled Yellow Fever Virus-specific CD8^+^ T cells(Minervina et al., 2020). We do note that the high degree of variability for S-specific CD8^+^ T cell recall in this cohort could mask subtle kinetic differences between primary and recall responses. Additionally, as the high naïve-precursor frequency of B7-N_105_ observed in our study and by others(Lineburg *et al*., 2021; Nguyen *et al*., 2021) may not apply to other pMHC-I specificities and may bias response kinetics, analysis of primary responses to other epitopes could be informative. Nevertheless, our data suggest that the protective benefit of circulating T cell memory may be more related to the maintenance of an expanded population of T cells with immediate effector function rather than accelerated activation or proliferation kinetics compared to a primary response. Given, however, that both the activation time and response peak time of primary N-specific CD8^+^ T cells in peripheral blood occur early after symptom onset, it is plausible that both primary and recalled responses contribute to viral clearance.

*In vitro* restimulation assays have typically been employed to enumerate vaccine-or infection-elicited T cells prior to BTI(Scurr *et al*., 2022; Tan et al., 2021a). However, the use of pMHC multimers allowed us to quantitively and qualitatively track epitope-specific T cells at markedly augmented resolution. For example, we noted a subset of individuals with no observable change in frequency of S-specific CD8^+^ T cells throughout BTI, but with robust evidence of activation. Restimulation-based T cell assays may be confounded during acute infection due to a combination of *in vivo* T cell activation and a lack of sensitivity to detect subtle changes in antigen-specific T cell frequency. Furthermore, our observation of a correlation between viral clearance and T cell recall was based on the measurement of activation, rather than T cell frequency. The ability to accurately determine the phenotype of antigen-specific CD8^+^ T cells directly *ex-vivo* has therefore provided critical insights into the recall of immunological memory and viral clearance that were not captured by simply enumerating such cells. A key consideration for future research will be to define how functional attributes of CD8^+^ T cells, like cytokine production or cytolytic potential, compare to phenotypic activation during BTI, which may guide the selection of optimal readouts for monitoring T cell recall and correlates of protection in larger cohorts.

We and others have previously described how cTFH responses serve as correlates of humoral immunity during viral infection and vaccination, including for SARS-CoV-2(Juno et al., 2020; Koutsakos et al., 2019; Koutsakos et al., 2018; Wragg *et al*., 2022). The low frequency of S-specific cTFH cells in our BTI cohort was therefore intriguing. It is possible that the interval between antigen exposures affects the emergence of cTFH, as second and third doses of mRNA-encoded spike also result in a limited induction of CXCR5^+^ S-specific CD4^+^ T cells relative to the first dose(Wragg *et al*., 2022). Similarly, analyses of influenza HA-specific CD4^+^ T cells after vaccination indicated limited induction of a cTFH response in individuals who were vaccinated within the past 12 months compared to those who were not(Wild et al., 2021). In addition to the limited S-specific cTFH phenotype, the lack of correlation between CD4^+^ T cell and antibody recall among our cohort was also surprising. This may reflect an overall less stringent involvement of T cell help in recall humoral responses, consistent with animal models(Hebeis et al., 2004; Zabel et al., 2017). Given evidence of germinal center activity following SARS-CoV-2 infection(Tan et al., 2022), further studies are thus needed to characterize lymphoid TFH and cTFH activity during recall following BTI with SARS-CoV-2 and its utility as a biomarker of humoral immunity in this context.

We note that our analysis is limited to specific HLA allotypes that may not be representative of the global population. Further studies are needed to establish equivalent pMHC multimer reagents across different HLA allotypes and antigens, as well as to develop strategies that may allow for their application in larger cohorts of genetically diverse populations. Although our study is focused on 26 individuals, the thorough kinetic analyses of 150 nasal swabs and 138 blood samples may facilitate more targeted sampling of larger cohorts at key timepoints defined in our study. Analysis of more diverse cohorts may facilitate further dissection of factors affecting immune recall and viral clearance including previous antigen exposures, age or disease severity, which was not possible in our cohort. Nonetheless, our study provides considerable evidence that T cell activation occurs before detectable expansion in the number of cells, and is correlated with virological control during BTI with SARS-CoV-2. This emphasizes the need to further understand the role of vaccine-elicited T cell immunity both mechanistically and as a correlate of protection.

## Acknowledgments

We thank the participants for their generous involvement and provision of samples. We thank molecular staff at the Victorian Infectious Diseases Reference Laboratory for performing RT-PCR, and Ms Grace Gare for technical assistance. We thank Dr. Julian Druce and Dr. Leon Caly at the Victorian Infectious Diseases Reference Laboratory for isolating and distributing SARS-CoV-2 virus isolates. We thank A/Prof Stuart Turville from the Kirby Institute, University of New South Wales for technical expertise and sharing the HAT-24 cell line. We acknowledge the Melbourne Cytometry Platform for provision of flow cytometry services. The work has been generously supported by the Morningside Foundation and by the Australian National Health and Medical Research Council grants 1149990, 1162760, 1194036 and 2004398; Australian Medical Research Future Fund grants 2005544, 2016062 and 2013870; The Victorian Government; and Australian National Health and Medical Research Council Investigator or Fellowship grants (M.K., A.K.W., J.A.J., H.-X.T., D.A.W., M.P.D., K.K, T.H.O.N. and S.J.K.).

## Author contributions

M.K. and J.A.J conceived and supervised the study. M.K., J.A.J, W.S.L., H.X.T. and A.K.W. designed experiments. M.K., J.A.J, W.S.L., G.T., P.K., K.C.L., T.T. performed experiments. J.N., T.A., H.E.K. and S.K. recruited, collected and processed participant samples. M.K., J.A.J, W.S.L., H.X.T., A.K.W., A.R., D.K., and M.P.D, contributed to data analysis. L.C.R, T.H.O.N., P.G.T, K.K., J.P, J.R. and D.A.W. contributed to methodology design. M.K., J.A.J, W.S.L., H.X.T., A.R., M.P.D and S.K. drafted the manuscript. All authors reviewed the final version of the manuscript.

## Figure legends

## Methods

### Study Participants

A cohort of previously vaccinated participants with a nasal PCR-confirmed breakthrough COVID-19 were recruited through contacts with the investigators and invited to provide serial blood samples following symptom onset (Table S1), some of whom were previously described in (Koutsakos *et al*., 2022). For all participants, whole blood was collected with sodium heparin anticoagulant. Plasma was collected and stored at -80°C, and PBMCs were isolated via Ficoll-Paque separation, cryopreserved in 10% DMSO/FCS and stored in liquid nitrogen. For some individuals, additional PBMC samples were available from participation in previous vaccine or SARS-CoV-2 infection studies(Juno *et al*., 2020; Wheatley et al., 2021; Wragg *et al*., 2022). Study protocols were approved by the University of Melbourne Human Research Ethics Committee (2021-21198-15398-3, 2056689), and all associated procedures were carried out in accordance with approved guidelines. All participants provided written informed consent in accordance with the Declaration of Helsinki. HLA typing was performed by the Victorian Transplantation and Immunogenetics Service.

### pHLA Production and Staining

Pro5 MHC Class I pentamers for HLA-A*02:01 (S_269-277_ YLQPRTFLL), HLA-A*03:01 (S_378-386_ KCYGVSPTK), and HLA-A*24:02 (S_1205-1213_ QYIKWPWYI) pre-conjugated to PE were purchased from ProImmune. ProT2 MHC Class II monomers for HLA-DRB1*15:01 (S_751-767_ NLLLQYGSFCTQLNRAL) were purchased from ProImmune and conjugated to Streptavidin-PE (BD Biosciences) at a molar ratio of 8:1 monomer:S-PE. HLA-B*07:02 (N_105-113_ SPRWYFYYL) and HLA-DP*04:01 (S_167-180_ TFEYVSQPFLMDLE) monomers were produced as described previously(Mudd *et al*., 2022; Nguyen *et al*., 2021). Monomers were conjugated to Streptavidin-APC at a molar ratio of 8:1 monomer:S-APC.

Cryopreserved PBMC were thawed in RPMI-1640 with 10% fetal calf serum (FCS) and penicillin-streptomycin (RF10), and washed. 7 – 13×10^6^ cells were washed in PBS with 2% FCS before incubation in 50nM dasatinib (Sigma) at 37°C for 30 minutes. HLA class II and HLA-B*07:02 tetramers were added directly to the tubes and stained at a final concentration of 2ug/mL for 60 minutes at 37°C. Class I pentamers were then added (final concentration 2ug/mL) and incubated for 15 minutes at room temperature in the dark. Cells were washed in PBS, stained with Live/Dead green (Life Technologies), and incubated for 30 minutes at 4°C with an antibody cocktail including: CD20 FITC (2H7), CD3 BUV805 (SK7), CD8 BUV496 (RPA-T8), CD71 BUV395 (M-A712), CD38 BUV737 (HB7, all from BD Biosciences), ICOS PerCP-Cy5.5 (C398.4A), CD4 APC-Cy7 (RPA-T4), CCR5 BV421 (J418F1), CXCR3 Pe-Dazzle (G02H57), PD-1 BV605 (EH12.2H7), CD45RA AlexaFluor700 (HI100, all from Biolegend), and CXCR5 PeCy7 (MU5UBEE; ThermoFisher). Cells were washed in PBS+2% FCS and permeabilised for 10 minutes in 70ul of Cytofix/Cytoperm (BD Biosciences). After washing in 2mL of PermWash, cells were incubated with anti-GzmB BV510 (GB11, BD Biosciences) for 30 minutes at 4°C. Cells were washed, resuspended in PBS+2% FCS, and acquired on a BD Fortessa using BD FACS Diva software.

### SARS-CoV-2 virus propagation and titration

Ancestral SARS-CoV-2 (VIC01) isolate was grown in Vero cells in serum-free DMEM with 1µg/ml TPCK trypsin while Omicron BA.1 and BA.2 strains were grown in Calu3 cells in DMEM with 2% FCS. Cell culture supernatants containing infectious virus were harvested on Day 3 for VIC01 and Day 4 for Omicron strains, clarified via centrifugation, filtered through a 0.45µM cellulose acetate filter and stored at -80°C. Infectivity of VIC01 stocks was determined by titration on Vero cells via cytopathic effect observation and calculated using the Reed-Muench method, as previously described(Juno *et al*., 2020). Infectivity of Omicron stocks was determined by titration on HAT-24 cells (a clone of transduced HEK293T cells stably expressing human ACE2 and TMPRSS2(Tea et al., 2021)). In a 96-well flat bottom plate, virus stocks were serially diluted five-fold (1:5-1:78,125) in DMEM with 5% FCS, added with 30,000 freshly trypsinised HAT-24 cells per well and incubated at 37°C. After 46 hours, 10µl of alamarBlue™ Cell Viability Reagent (ThermoFisher) was added into each well and incubated at 37°C for 1 hour. The reaction was then stopped with 1% SDS and read on a FLUOstar Omega plate reader (excitation wavelength 560nm, emission wavelength 590nm). The relative fluorescent units (RFU) measured were used to calculate %viability (‘sample’ ÷ ’no virus control’ × 100), which was then plotted as a sigmoidal dose response curve on Graphpad Prism to obtain the virus dilution that induces 50% cell death (50% lethal infectious dose; LD_50_). Each virus was titrated in quintuplicate in three independent experiments to obtain mean LD_50_ values.

### SARSCoV-2 microneutralization assay with ELISA-based readout

For Delta breakthrough infections, plasma neutralization activity against ancestral SARS-CoV-2 (CoV/Australia/VIC/01/2020 strain) was measured using a microneutralization assay as previously described(Koutsakos *et al*., 2022). 96-well flat bottom plates were seeded with Vero cells (20,000 cells per well in 100µl). The next day, Vero cells were washed once with 200 µl serum-free DMEM and added with 150µl of infection media (serum-free DMEM with 1.33 µg/ml TPCK trypsin). 2.5-fold serial dilutions of heat-inactivated plasma (1:20-1:12207) were incubated with SARS-CoV-2 virus at 2000 TCID_50_/ml at 37°C for 1 hour. Next, plasma-virus mixtures (50µl) were added to Vero cells in duplicate and incubated at 37°C for 48 hours. ‘Cells only’ and ‘virus+cells’ controls were included to represent 0% and 100% infectivity respectively. After 48 hours, all cell culture media were carefully removed from wells and 200 µl of 4% formaldehyde was added to fix the cells for 30 mins at room temperature. The plates were then dunked in a 1% formaldehyde bath for 30 minutes to inactivate any residual virus prior to removal from the BSL3 facility. Cells were washed once in PBS and then permeabilized with 150µl of 0.1% Triton-X for 15 minutes. Following one wash in PBS, wells were blocked with 200µl of blocking solution (4% BSA with 0.1% Tween-20) for 1 hour. After three washes in PBST (PBS with 0.05% Tween-20), wells were incubated with 100µl of rabbit polyclonal anti-SARS-CoV N antibody (Rockland, #200-401-A50) at a 1:8000 dilution in dilution buffer (PBS with 0.2% Tween-20, 0.1% BSA and 0.5% NP-40) for 1 hour. Plates were then washed six times in PBST and added with 100µl of goat anti-rabbit IgG (Abcam, #ab6721) at a 1:8000 dilution for 1 hour. After six washes in PBST, plates were developed with TMB and stopped with 0.15M H_2_SO_4_. OD values read at 450nm were then used to calculate %neutralization with the following formula: (‘Virus + cells’ – ‘sample’) ÷ (‘Virus + cells’ – ‘Cells only’) × 100. IC_50_ values were determined using four-parameter nonlinear regression in GraphPad Prism with curve fits constrained to have a minimum of 0% and maximum of 100% neutralization.

### SARS-CoV-2 microneutralisation assay with viability dye readout

For Omicron breakthrough infections, plasma neutralization activity against Omicron BA.1 and BA.2 was measured in HAT-24 cells using a viability dye readout. In 96-well flat bottom plates, heat-inactivated plasma samples were diluted 2.5-fold (1:20-1:12,207) in duplicate and incubated with SARS-CoV-2 virus at a final concentration of 2× LD50 at 37°C for 1 hour. Next, 30,000 freshly trypsinised HAT-24 cells in DMEM with 5% FCS were added and incubated at 37°C. ‘Cells only’ and ‘Virus+Cells’ controls were included to represent 0% and 100% infectivity respectively. After 46 hours, 10µl of alamarBlue™ Cell Viability Reagent (ThermoFisher) was added into each well and incubated at 37°C for 1 hour. The reaction was then stopped with 1% SDS and read on a FLUOstar Omega plate reader (excitation wavelength 560nm, emission wavelength 590nm). The relative fluorescent units (RFU) measured were used to calculate %neutralisation with the following formula: (‘Sample’ – ‘Virus+Cells’) ÷ (‘Cells only’ – ‘Virus+Cells’) × 100. IC50 values were determined using four-parameter non-linear regression in GraphPad Prism with curve fits constrained to have a minimum of 0% and maximum of 100% neutralisation.

### Analysis of viral RNA load by qPCR

For viral RNA extraction, 200 μL of nasal swab sample was extracted with the QIAamp 96 Virus QIAcube HT kit (Qiagen, Germany) on the QIAcube HT System (Qiagen) according to manufacturer’s instructions. Purified nucleic acid was then immediately converted to cDNA by reverse transcription with random hexamers using the SensiFAST cDNA Synthesis Kit (Bioline Reagents, UK) according to manufacturer’s instructions. cDNA was used immediately in the rRT-PCR or stored at -20oC. Three microlitres of cDNA was added to a commercial real-time PCR master mix (PrecisionFast qPCR Master Mix; Primer Design, UK) in a 20 μL reaction mix containing primers and probe with a final concentration of 0.8µM and 0.1µM for each primer and the probe, respectively. Samples were tested for the presence of SARS-CoV-2 nucleocapsid (N) genes using previously described primers and probes (Chan et al., 2020; Corman et al., 2020). Thermal cycling and rRT-PCR analyses for all assays were performed on the ABI 7500 FAST real-time PCR system (Applied Biosystems, USA) with the following thermal cycling profile: 95oC for 2 min, followed by 45 PCR cycles of 95oC for 5 s and 60oC for 30 s for N gene.

### Modelling of viral and immune kinetics

We used a piecewise model to estimate the activation time and growth rate of various immune responses following breakthrough infections. The model of the immune response *y* for subject *i* at time *y*_!_ can be written as:

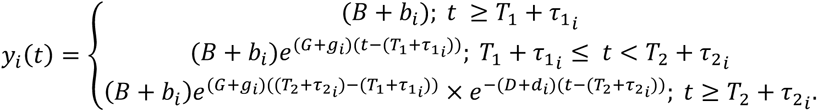

The model has 5 parameters; *B*, *G*, *T*_1_, *D*, and *T*_2_. For a period before *T*_1_, we assumed a constant baseline value *B* for the immune response. After the activation time *T*_2_, the immune response will grow at a rate of *G* until *T*_2_. From *T*_2_, the immune response will decay at a rate of *D*. For each subject *i*, the parameters were taken from a normal distribution, with each parameter having its own mean (fixed effect). A diagonal random effect structure was used, where we assumed there was no correlation within the random effects. The model was fitted to the log-transformed data values, with a constant error model distributed around zero with a standard deviation σ. Model fitting was performed using MonolixR2019b. A binary covariate was used to quantify the difference in parameters between different groups (i.e. S-specific CD8 vs N-specific CD8 responses), and significance was determined based on the value of this binary covariate using a Wald test. Spearman correlation analyses were conducted based on the estimated parameters for each individual in GraphPad Prism 9.

## Supplementary material

**Supplementary Table 1.** Cohort demographics

| Subject | Gender | Age | # Prior Vaccine doses | Last vaccine to symptom onset (days) | Class I Alleles (relevant to this study) | Class II Alleles (relevant to this study) | VOC |
| --- | --- | --- | --- | --- | --- | --- | --- |
| CP105 | M | 29 | 2 | N.D. | A*02:01 | -- | Delta |
| CP106 | F | 30 | 2 | 139 | -- | DP*04:01 | Delta |
| CP107 | F | 31 | 2 | 77-106 | A*02:01 | DP*04:01 | Delta |
| CP108 | M | 29 | 2 | 132 | -- | DP*04:01 | Delta |
| COR034 | M | 31 | 2 | 126 | A*02:01, B*07:02 | DR*15:01, DP*04:01 | Delta |
| COR136 | F | 50 | 2 | 158 | A*02:01, A*03:01 | DP*04:01 | Delta |
| COR015 | F | 27 | 3 | 39 | A*02:01 | -- | BA.1 |
| COR032 | F | 23 | 3 | 33 | A*24:02, B*07:02 | DR*15:01, DP*04:01 | BA.1 |
| CP110 | M | 24 | 2 | 84 | A*02:01 | DP*04:01 | BA.1 |
| CP111 | M | 34 | 2 | 83 | A*24:02 | -- | BA.1 |
| CP112 | F | 35 | 2 | 90 | A*02:01<sup>^</sup> | DP*04:01 | BA.1 |
| COR198 | F | 60 | 3 | 39 | A*02:01 | DP*04:02 | BA.1 |
| CP69* | F | 36 | 2 | 111 | A*02:01 | DR*15:01, DP*04:01 | BA.1 |
| CP40* | M | 63 | 3 | 49 | -- | DR*15:01, DP*04:01 | BA.2 |
| COR291 | M | 50 | 3 | 134 | A*02:01 | DR*15:01 | BA.2 |
| COR043 | F | 55 | 3 | 64 | A*02:01 | DP*04:01 | BA.2 |
| CP117 | M | 40 | 3 | 52 | A*03:01, B*07:02 | DR*15:01, DP*04:01 | BA.2 |
| COR281 | F | 43 | 3 | 129 | A*03:01, B*07:02 | DR*15:01, DP*04:01 | BA.2 |
| CP118 | M | 66 | 3 | 100 | -- | DP*04:01 | BA.2 |
| COR274 | F | 57 | 3 | 155 | A*03:01 | DP*04:01, 04:02 | BA.2 |
| COR275 | M | 62 | 3 | 83 | A*02:01, B*07:02 | DR*15:01, DP*04:01 | BA.2 |
| COR215 | F | 55 | 3 | 125 | A*24:02, B*07:02 | DR*15:01, DP*04:01 | BA.2 |
| COR039 | F | 58 | 3 | 169 | A*24:02, B*07:02 | DR*15:01, DP*04:01 | BA.2 |

\*Previously infected during Hu-1 wave. <sup>^</sup>Data not shown, due to lack of A\*02:01 pentamer binding in any longitudinal samples from participant; N.D., not determined.

**Supplementary Table 2.** Piecewise linear regression parameters of viral kinetics and neutralising antibodies (with 95% CI). Values in bold indicate a significant difference between VOCs.

|  |  | Initial value | Delay<br>(days PSO) | Growth Rate<br>(per day) | Peak Time<br>(days PSO) | Decay Rate<br>(per day) |
| --- | --- | --- | --- | --- | --- | --- |
| Viral<br>RNA<br>(ΔCt) | Delta | 36.5<br>(44.71 – 28.29) | N/A | 5.47<br>(2.83 – 10.57) | 3.22<br>(2.53 – 4.1) | **3.97**<br>(3.03 – 5.22) |
|  | BA.1 | 31.79<br>(49.98 – 13.6) | N/A | 4.41<br>(0.82 – 23.77) | 2.43<br>(1.24 – 4.78)* | **2.13**<br>(1.03 – 4.43) |
|  | BA.2 | 24.2<br>(49.5 – 1.1) | N/A | 6.73<br>(0.32 – 139.23) | 1.12<br>(0.47 – 2.63) | **2.04**<br>(1.12 – 3.73) |
|  | Nab (IC<sub>50</sub>) | 51.29<br>(30.52 – 86.18) | 4.3<br>(2.2 – 8.42) | 0.12<br>(0.084 – 0.18) | 14.58<br>(12.23 – 17.39) | N/A |

**Supplementary Table 3.** Estimates of spike-specific CD4^+^ T cell expansion and activation. Pooled estimates from both epitopes are shown as the epitope-specific estimates were not significantly different for any parameter.

|  | Initial (%) | Delay<br>(days PSO) | Growth Rate<br>(per day) | Peak Time<br>(days PSO) | Decay Rate<br>(per day) |
| --- | --- | --- | --- | --- | --- |
| %pMCH<sup>+</sup> of<br>CD4<sup>+</sup> | 0.0093<br>(0.0071 – 0.012) | 2.51<br>(1.86 – 3.4) | 0.24<br>(0.16 – 0.35) | 5.42<br>(4.81 – 6.1) | 0.014<br>(0.01 – 0.019) |
| %CD38<sup>+</sup><br>ICOS<sup>+</sup> of<br>pMCH<sup>+</sup> | 2.44<br>(1.1 – 5.42) | 1.12<br>(0.7 – 1.8) | 0.62<br>(0.5 – 0.76) | 3.60<br>(3.22 – 4.01) | 0.057<br>(0.051 – 0.063) |

**Supplementary Table 4.** Estimates of spike-specific CD4^+^ T cell phenotypic parameters (with 95% CI). Pooled estimates from both epitopes are shown if the epitope-specific estimates were not significantly different for any parameter. Separate epitope-specific estimates are shown if at least one of the parameters of that makers were significantly different between epitopes. Values in bold indicate a significant difference between epitopes for the indicated marker.

|  | Initial (%) | Growth Rate (per<br>day) | Peak Time (days<br>PSO) | Decay Rate<br>(per day) |
| --- | --- | --- | --- | --- |
| %CCR5<sup>+</sup><br>(pooled epitopes) | 16.59<br>(12 – 22.96) | 0.26<br>(0.19 0 0.35) | 2.48<br>(2.15 – 2.85) | 0.0064<br>(0.0043 – 0.0095) |
| %CXCR3<sup>+</sup><br>(pooled epitopes) | 14.4<br>(7.51 – 27.8) | 0.27<br>(0.13 – 0.54) | 1.93<br>(1.25 – 2.96) | 0.0064<br>(0.0032 – 0.013) |
| %GzmB<sup>+</sup><br>(pooled epitopes) | 7.34<br>(4.27 – 12.62) | 0.13<br>(0.04 – 0.4) | 1.97<br>(0.88 – 4.38) | 0.019<br>(0.013 – 0.029) |
| %PD-1<sup>+</sup> (DP04) | 22.38<br>(10.49 – 47.78) | 0.11<br>(0.05 – 0.26) | 4.05<br>(3.29 – 5) | **0.0078**<br>(0.004 – 0.015) |
| %PD-1<sup>+</sup> (DR15) | 34.04<br>(6.7 – 172.82) | 0.12<br>(0.019 – 0.78) | 2.71<br>(1.46 – 5.03) | **0.00047**<br>(0.00023 – 0.0093) |
| %CD71<sup>+</sup> (DP04) | 1.05<br>(0.39 – 2.86) | 0.38<br>(0.23 – 0.62) | 3.35<br>(2.73 – 4.11) | **0.077**<br>(0.056 – 0.11) |
| %CD71<sup>+</sup> (DR15) | 1.32 (0.08 – 21.71) | 0.45 (0.00011 –<br>1.821) | 1.64 (0.001 – 2.588) | **0.038**<br>(0.014 – 0.1) |

**Supplementary Table 5.**
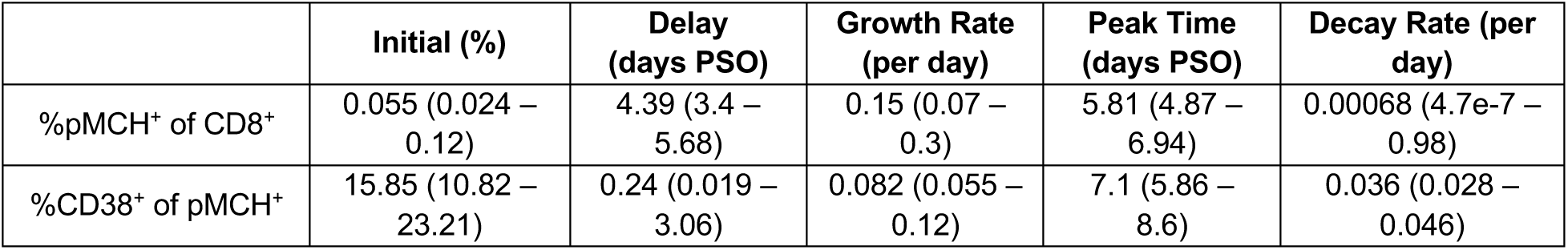
Estimates of S-specific CD8^+^ T cell expansion and activation (with 95% CI). Pooled estimates from all three epitopes are shown as the epitope-specific estimates were not significantly different for any parameter.

|  | Initial (%) | Delay (days PSO) | Growth Rate (per day) | Peak Time (days PSO) | Decay Rate (per day) |
| --- | --- | --- | --- | --- | --- |
| %pMCH<sup>+</sup> of CD8<sup>+</sup> | 0.055 (0.024 – 0.12) | 4.39 (3.4 – 5.68) | 0.15 (0.07 – 0.3) | 5.81 (4.87 – 6.94) | 0.00068 (4.7e-7 – 0.98) |
| %CD38<sup>+</sup> of pMCH<sup>+</sup> | 15.85 (10.82 – 23.21) | 0.24 (0.019 – 3.06) | 0.082 (0.055 – 0.12) | 7.1 (5.86 – 8.6) | 0.036 (0.028 – 0.046) |

**Supplementary Table 6.** Estimates of S-specific CD8^+^ T cell phenotypic parameters (with 95% CI). Pooled estimates from all three epitopes are shown.

|  | Initial (%) | Growth Rate (per day) | Peak Time (days PSO) | Decay Rate (per day) |
| --- | --- | --- | --- | --- |
| %CCR5<sup>+</sup> | 40.74 (34.41 – 48.23) | 0.031 (0.018 – 0.052) | 6.89 (5.17 – 9.17) | 0.0032 (0.0015 – 0.0068) |
| %CXCR3<sup>+</sup> | 6.68 (2.93 – 15.26) | 0.2 (0.12 – 0.34) | 3.49 (2.56 – 4.75) | 0.0093 (0.0049 – 0.017) |
| %GzmB<sup>+</sup> | 23.99 (13.77 – 41.79) | 0.12 (0.007 – 2.14) | 1.4 (0.15 – 12.58) | 0.0069 (0.0039 – 0.012) |
| %CD71<sup>+</sup> | 0.64 (0.2 – 2) | 0.23 (0.14 – 0.38) | 5.36 (4.25 – 6.77) | 0.033 (0.022 – 0.049) |
| %PD-1<sup>+</sup> | 7.52 (4.53 – 12.46) | 0.096 (0.04 – 0.23) | 3.32 (2.14 – 5.16) | 0.0079 (0.0047 – 0.013) |

**Supplementary Table 7.** Estimates of S- and N-specific CD8^+^ T cell expansion and activation (with 95% CI). Values in bold indicate a significant difference between epitopes for the indicated marker.

|  | Initial (%) | Delay (days PSO) | Growth Rate (per day) | Peak Time (days PSO) | Decay Rate (per day) |
| --- | --- | --- | --- | --- | --- |
| %pMCH<sup>+</sup> of CD8<sup>+</sup> (S) | **0.098**<br>(0.012 – 0.82) | 6.29<br>(1.49 – 26.46) | 0.087<br>(0.013 – 0.556) | 7.69<br>(3.56 – 16.57) | 0 (NA) |
| %pMCH<sup>+</sup> of CD8<sup>+</sup> (N) | **0.012**<br>(0.0048 – 0.03) | 3.06<br>(1.62 – 5.81) | 0.12<br>(0.071 – 0.2) | 8.17<br>(6.32 – 10.56) | 0 (NA) |
| %CD38<sup>+</sup> (S) | 10.16<br>(1.66 – 62.36) | 0.95<br>(0.001 – 86.85) | 0.073 (0.01 – 0.52) | 8.29<br>(4.45 – 15.43) | **0.047**<br>(0.014 – 0.16) |
| %CD38<sup>+</sup> (N) | 17.78<br>(9.58 – 33) | 3.63<br>(2.28 – 5.79) | 0.19<br>(0.077 – 0.45) | 6.89<br>(5.26 – 9.02) | **0.019**<br>(0.011 – 0.034) |

**Supplementary Table 8.**
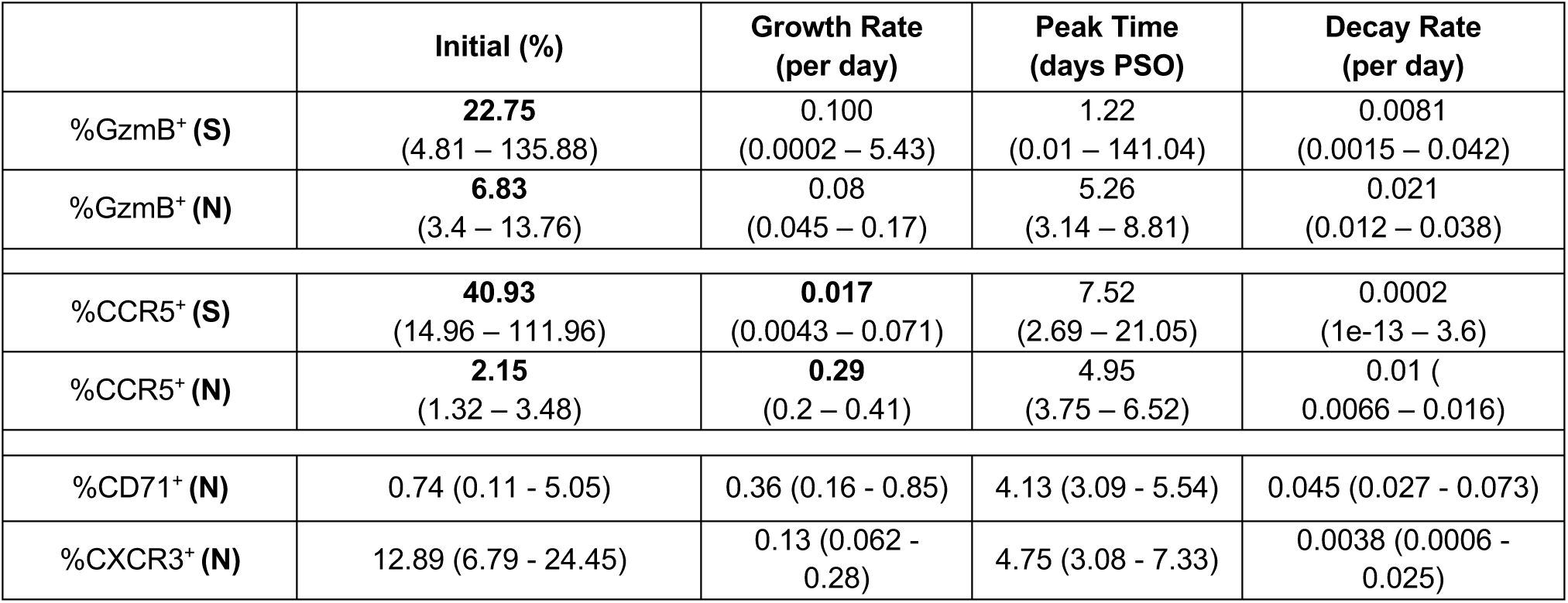
Estimates of S- and N-specific CD8^+^ T cell phenotypic parameters (with 95% CI) for individuals presented in Figure 4. Estimates for N-specific CD8^+^ T cells only are shown if the S -and N-specific estimates were not significantly different for any parameter (estimates for S only are shown in Supplementary Table 6. Separate epitope-specific estimates are shown if at least one of the parameters of that makers were significantly different between epitopes. Values in bold indicate a significant difference between epitopes for the indicated marker. Growth and decay parameters for PD-1 expression kinetics could not be determined.

|  | **Initial (%)** | **Growth Rate**<br>**(per day)** | **Peak Time**<br>**(days PSO)** | **Decay Rate**<br>**(per day)** |
| --- | --- | --- | --- | --- |
| %GzmB<sup>+</sup> (**S**) | **22.75**<br>(4.81 – 135.88) | 0.100<br>(0.0002 – 5.43) | 1.22<br>(0.01 – 141.04) | 0.0081<br>(0.0015 – 0.042) |
| %GzmB<sup>+</sup> (**N**) | **6.83**<br>(3.4 – 13.76) | 0.08<br>(0.045 – 0.17) | 5.26<br>(3.14 – 8.81) | 0.021<br>(0.012 – 0.038) |
| %CCR5<sup>+</sup> (**S**) | **40.93**<br>(14.96 – 111.96) | **0.017**<br>(0.0043 – 0.071) | 7.52<br>(2.69 – 21.05) | 0.0002<br>(1e-13 – 3.6) |
| %CCR5<sup>+</sup> (**N**) | **2.15**<br>(1.32 – 3.48) | **0.29**<br>(0.2 – 0.41) | 4.95<br>(3.75 – 6.52) | 0.01<br>(0.0066 – 0.016) |
| %CD71<sup>+</sup> (**N**) | 0.74 (0.11 - 5.05) | 0.36 (0.16 - 0.85) | 4.13 (3.09 - 5.54) | 0.045 (0.027 - 0.073) |
| %CXCR3<sup>+</sup> (**N**) | 12.89 (6.79 - 24.45) | 0.13 (0.062 - 0.28) | 4.75 (3.08 - 7.33) | 0.0038 (0.0006 - 0.025) |

## Supplementary Figure legends

**Figure S1.**
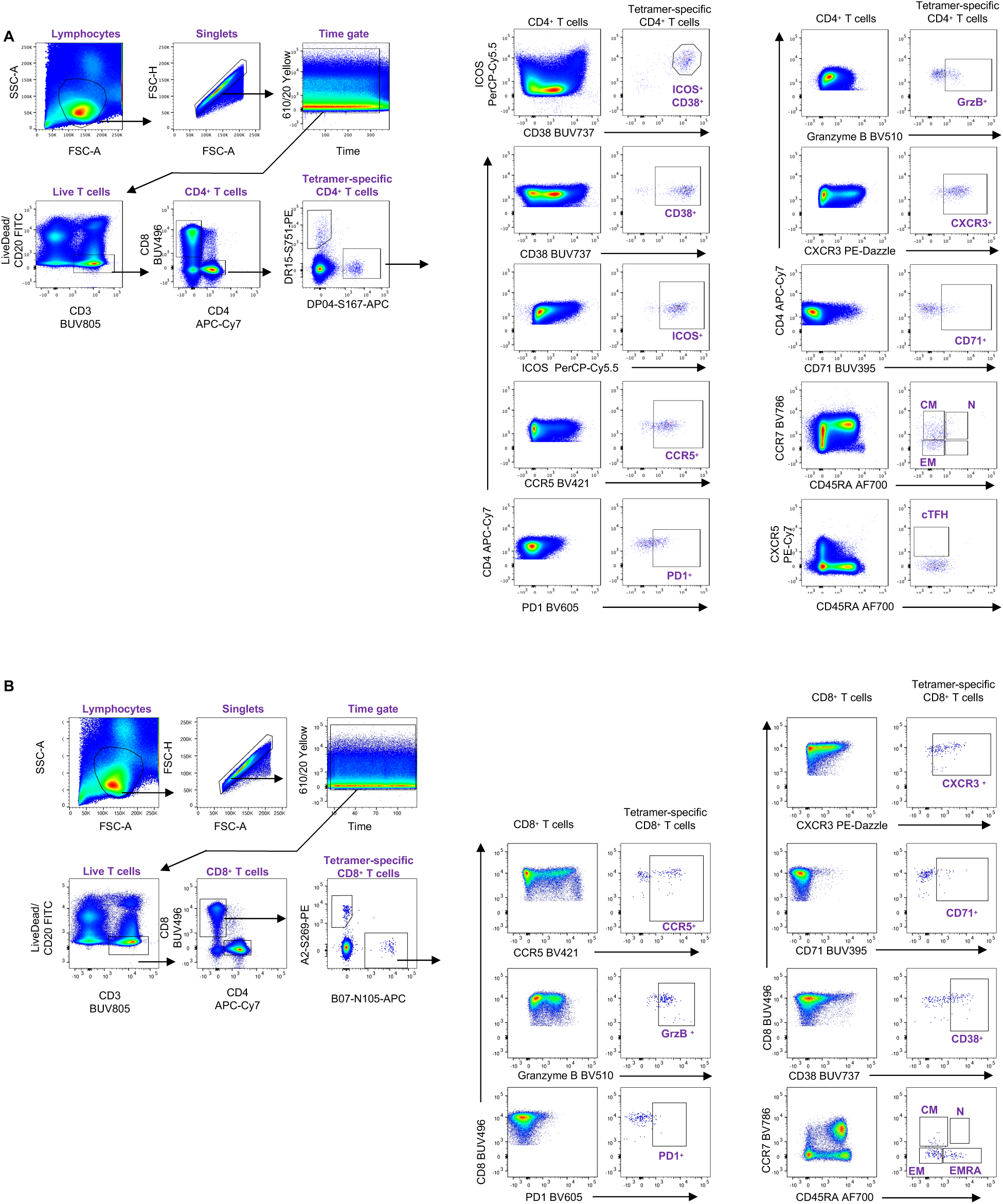
Gating strategy for the identification of pMHC-specific CD4^+^ and CD8^+^ T cells and their phenotypic characterisation. **(A)** Lymphocytes were identified by FSC-A vs SSC-A gating, followed by doublet exclusion (FSC-A vs FSC-H), a time gate and then gating on live T cells (CD3^+^CD19^-^) and subsequently CD4^+^CD8^-^ cells. HLA-DR*15-S_751_ and HLA-DP*04-S_167_-specific cells were identified within CD4^+^ T cells, and phenotyped as indicated, with the total CD4^+^ T cell population serving as a reference for gating of phenotypic markers. **(B)** CD8^+^ T cell were identified as CD4^-^CD8^+^ within live T cells gated as in (A). HLA-A*02-S_269_, HLA-A*03-S_378_ and HLA-A*24-S_1208_ and HLA-B*07-N_105_-specific CD8^+^ T cells were identified within CD8^+^ T cells and phenotyped as indicated, with the total CD8^+^ T cell population serving as a reference for gating of phenotypic markers.

**Figure S2.**
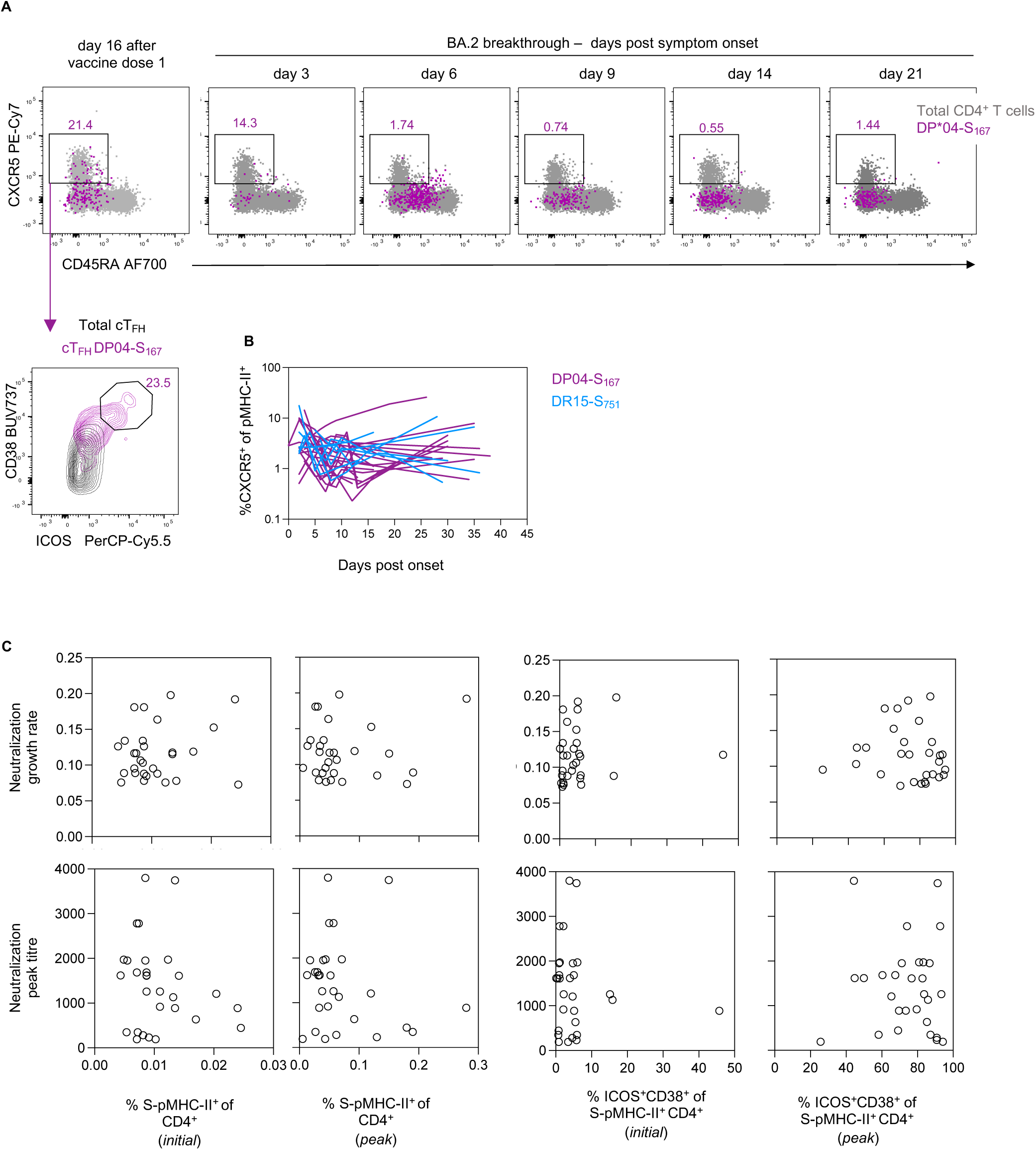
Limited cTFH phenotype following BTI. **(A)** Representative flow cytometry plots of the cTFH phenotype (CXCR5^+^CD45RA^-^) for HLA-DP*04-S_167_-specific CD4^+^ T cells. A post-vaccination sample was included in acquisition and analysis, serving as a reference for the cTFH activation and cTFH activation (CD38/ICOS expression). **(B)** Kinetics of CXCR5^+^ cells for both pMHC-II populations, n=19 for DP*04-S_167_ and n=9 for DR*15-S_751_. **(C)** Correlations between the initial or peak frequency of S-pMHC-II^+^ CD4^+^ T cells and initial or peak frequency of ICOS^+^CD38^+^ S-pMHC-II^+^ CD4^+^ T cells with the growth rate and peak value of neutralising antibody titres. Spearman correlation coefficient and p-values along with a linear regression line are shown for statistically significant comparisons (p<0.05), n=29 datapoints, pooled for both S-pMHC-II^+^ CD4^+^ T cell populations.

**Figure S3.**
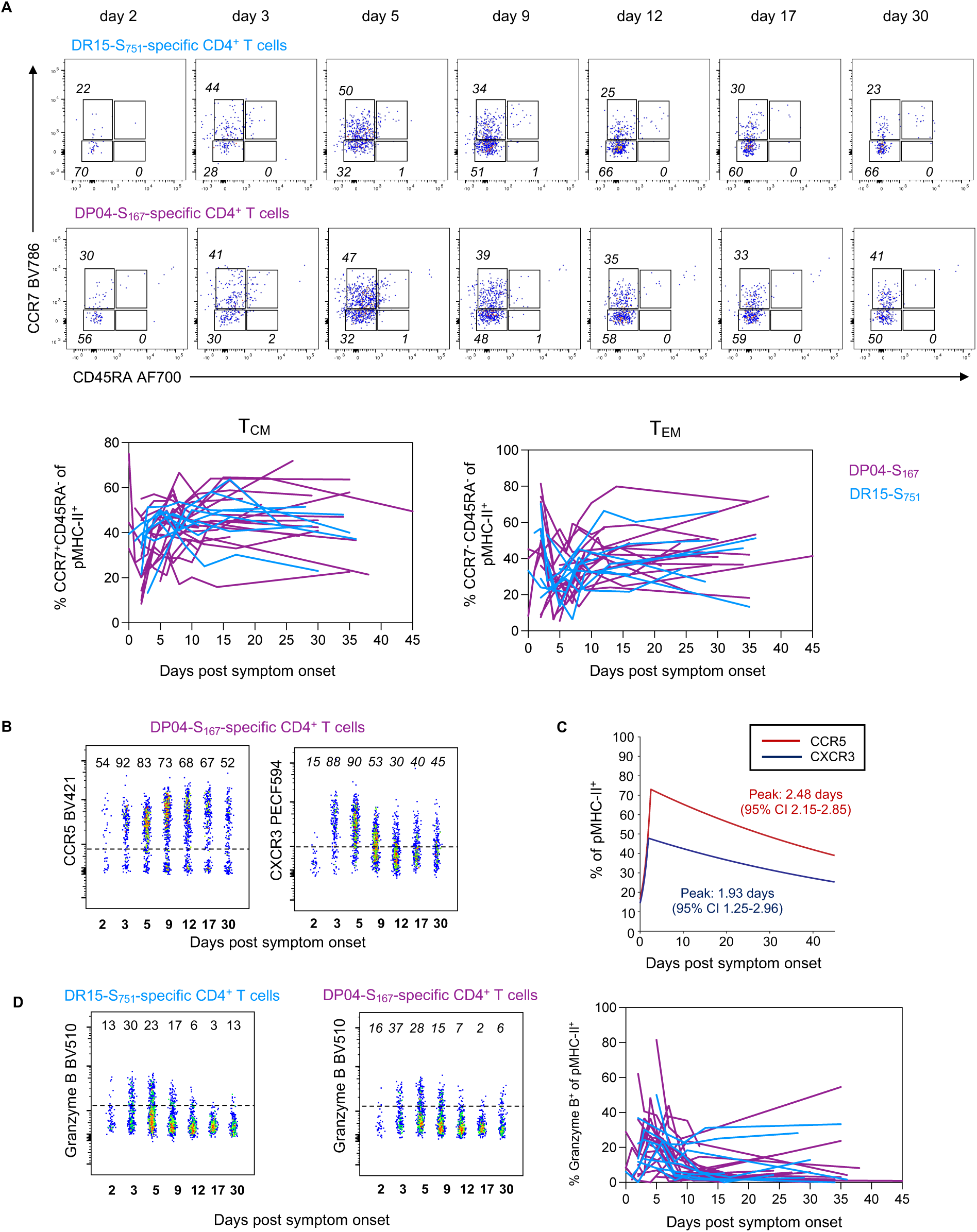
Phenotype of S-specific CD4^+^ T cells. **(A)** Representative flow cytometry plots of HLA-DR*15-S_751_ and HLA-DP*04-S_167_-specific CD4^+^ T cells from a single participant showing co-expression of CCR7 and CD45RA. Kinetics of T_CM_ and T_EM_ for both pMHC-II populations. **(B)** Flow cytometry plots of HLA-DP*04-S_167_-specific CD4^+^ T cells from a single participant for CCR5 and CXCR5 expression, representative of the data shown in Fig 2. **(C)** Estimated kinetics of CCR5 and CXCR3 expression. The lines indicate the mean estimate for measurement from the piecewise linear regression model, using pooled data from both pMHC-II populations as no significant differences were found between the two. **(D)** Flow cytometry plots of HLA-DR*15-S_751_ and HLA-DP*04-S_167_-specific CD4^+^ T cells from a single participant showing expression of Granzyme B, with kinetics from all participants shown both pMHC-II populations. Throughout the figure, coloured lines represent individual donors for each pMHC-specific population, n=19 for DP*04-S_167_ and n=9 for DR*15-S_751_.

**Figure S4.**
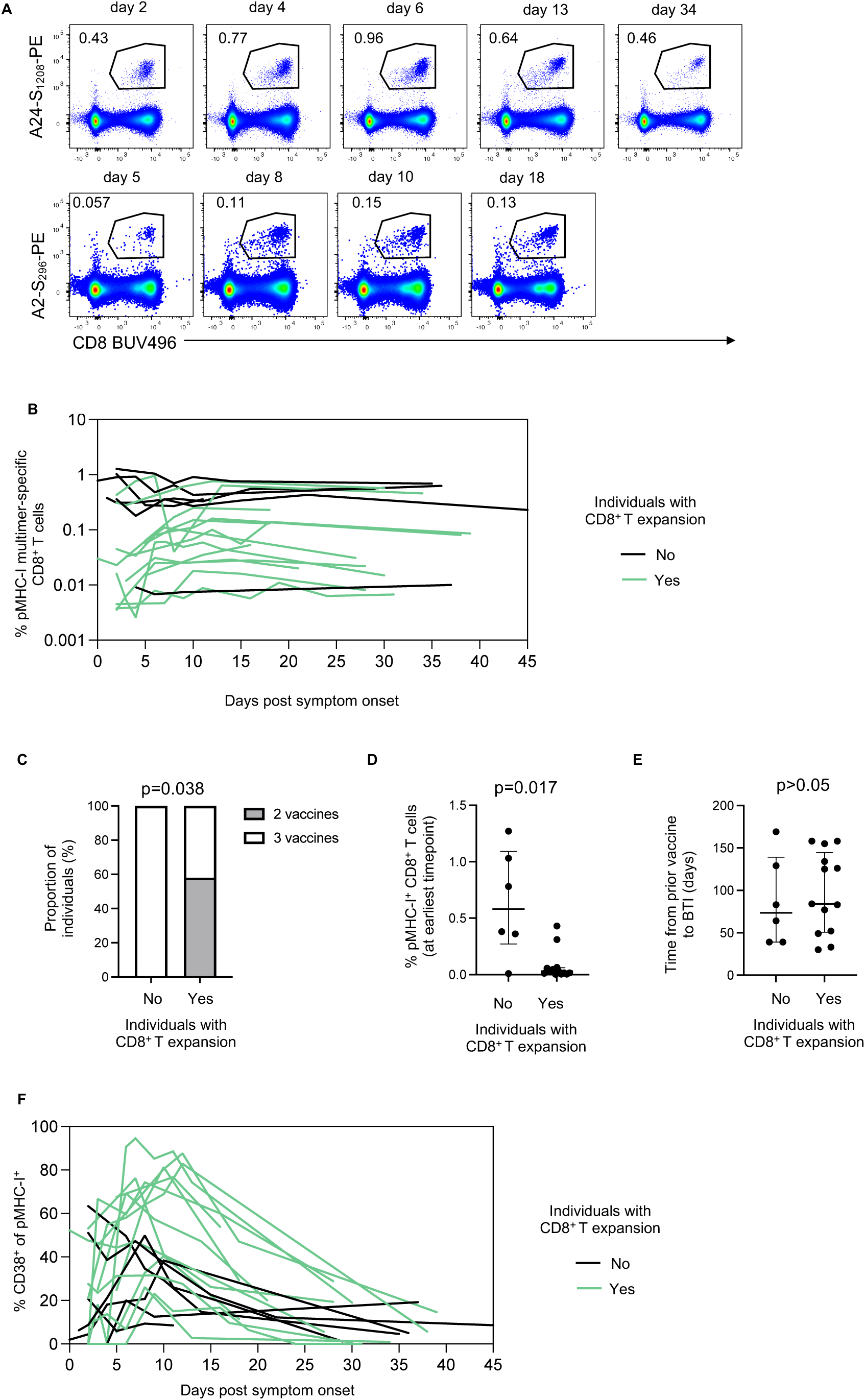
CD8^+^ T cell responses following BTI. **(A)** Representative flow cytometry plots of HLA-A*02-S_269_ and HLA-A*24-S_1208_ -specific CD8^+^ T cell kinetics. **(B)** pMHC-I^+^ CD8^+^ T cell kinetics colour coded for individuals with an observable expansion and those without. **(C)** Vaccination history of individuals with an observable expansion and those without. **(D)** pMHC-I^+^ CD8^+^ T cell frequency at earliest available timepoint for individuals with an observable expansion and those without. **(E)** Time from last vaccination to BTI for individuals with an observable and those without. **(F)** CD38^+^ phenotype for pMHC-I^+^ CD8^+^ T cell kinetics colour coded for individuals with an observable expansion and those without.

**Figure S5.**
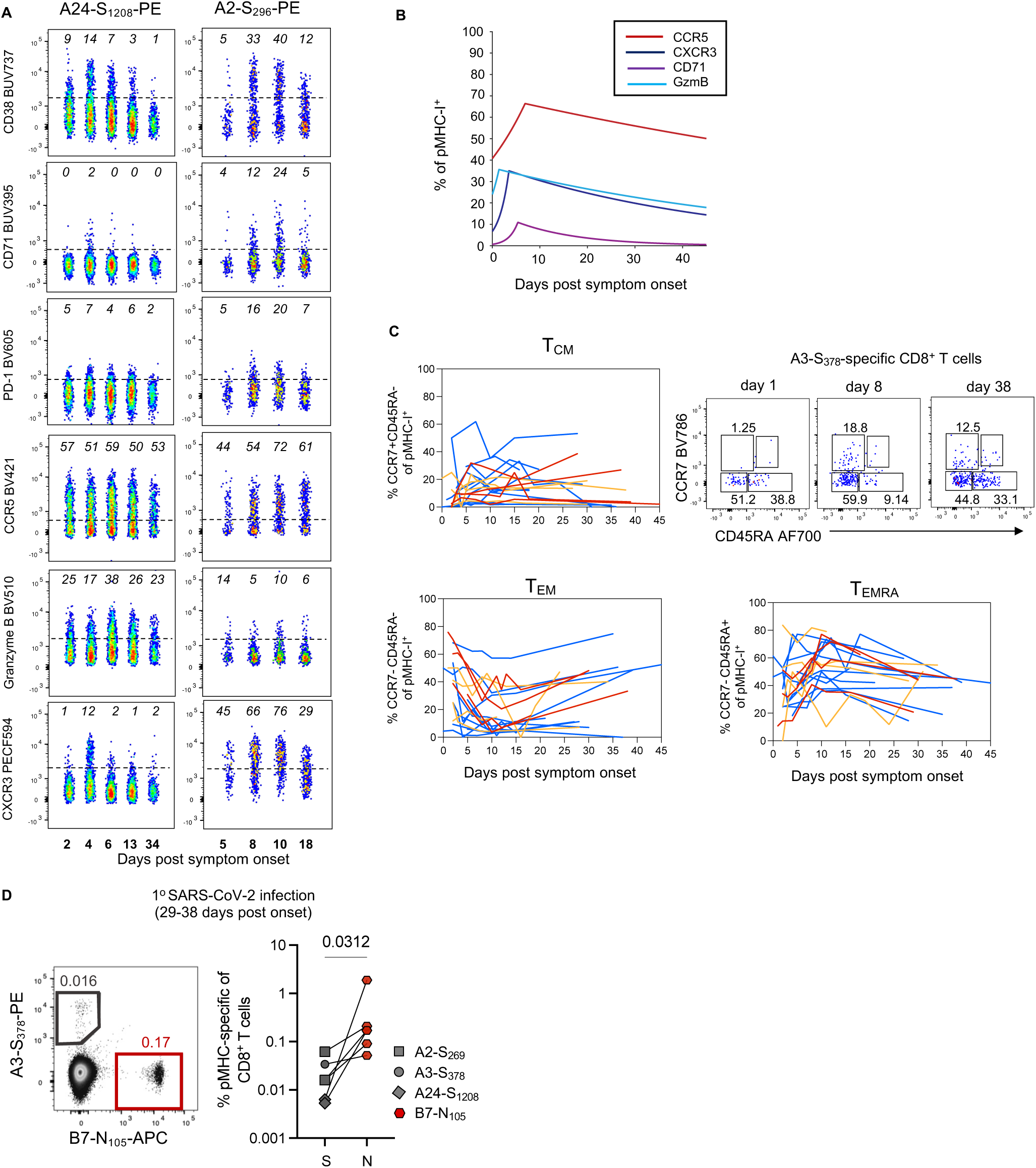
Phenotype of S-specific CD8^+^ T cells. **(A)** Flow cytometry plots of HLA-A*02-S_269_ and HLA-A*24-S_1208_-specific CD8^+^ T cells from a single participant for different phenotypic markers, representative of the data shown in Fig 3. **(B)** Estimated kinetics of CCR5, CXCR3, CD71 and GzmB expression. The lines indicate the mean estimate for measurement from the piecewise linear regression model, using pooled data from both pMHC-I populations as no significant differences were found between the two. **(C)** Representative flow cytometry plots of HLA-A*03-S_378_ -specific CD8^+^ T cells from a single participant showing co-expression of CCR7 and CD45RA. Kinetics of T_CM_ T_EM_ and T_EMRA_ for all 3 pMHC-I populations. Throughout the figure, coloured lines represent individual donors for each pMHC-I-specific population, n=11 for A*02-S_269_, n=4 A*03-S_378_ and n=4 for A*24-S_1208_. **(D)** Frequency of spike-specific and nucleoprotein-specific CD8^+^ T cells in convalescent samples from primary SARS-CoV-2 infection. For (C), n=6 donors with paired analysis of spike-specific CD8^+^ T cells (either A*02, A*03 and A*24) and B*07-N_105._

**Figure S6.**
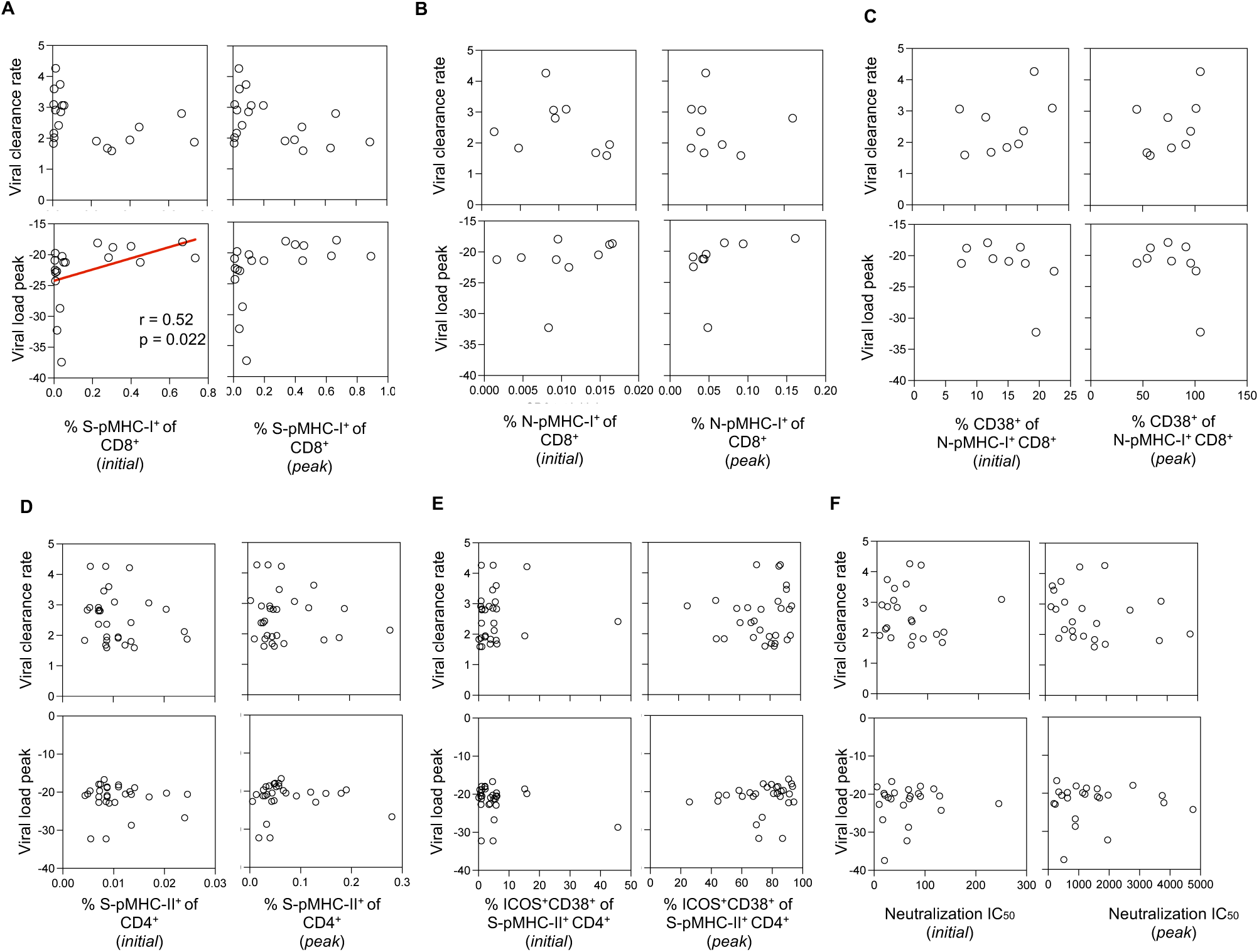
Correlations between immune recall and viral clearance. Correlations between the initial or peak frequency of **(A)** S-pMHC-I^+^ CD8^+^ T cells, **(B)** N-pMHC-I^+^ CD8^+^ T cells or **(C)** CD38^+^ N-pMHC-I^+^ CD8^+^ T cells, **(D)** S-pMHC-II^+^ CD4^+^ T cells, **(E)** ICOS^+^CD38^+^S-pMHC-II^+^ CD4^+^ T cells, **(F)** neutralisation IC_50_ titre against infecting or antigenically similar live virus and viral clearance rate or peak Ct value (amongst available timepoints). Throughout the figure Spearman correlation coefficient and p-values along with a linear regression line are shown for statistically significant comparisons (p<0.05), n=19 for (A), n=9 for (B-C), n=29 datapoints, pooled for both S-pMHC-II^+^ CD4^+^ T cell populations for (D-E), n=23 for (F).

